# Improving American chestnut resistance to two invasive pathogens through genome-enabled breeding

**DOI:** 10.1101/2025.01.30.635736

**Authors:** Jared W. Westbrook, Joanna Malukiewicz, Qian Zhang, Avinash Sreedasyam, Jerry W. Jenkins, Vasiliy Lakoba, Sara Fitzsimmons, Jamie Van Clief, Kendra Collins, Stephen Hoy, Cassie Stark, Lake Grabowski, Eric Jenkins, Thomas M. Saielli, Benjamin T. Jarrett, Lucinda J. Wigfield, Lauren M. Kerwien, Ciera Wilbur, Alexander M. Sandercock, J. Hill Craddock, Paola Zannini, Susanna Keriö, Tetyana Zhebentyayeva, Shenghua Fan, Austin M. Thomas, Albert G. Abbott, C. Dana Nelson, Xiaoxia Xia, Melissa Williams, LoriBeth Boston, Christopher Plott, Florian Carle, Jack Swatt, Jack Ostroff, Steven N. Jeffers, Kathleen Mckeever, Erica Smith, Thomas J. Ellis, Joseph B. James, Paul Sisco, Andrew Newhouse, Erik Carlson, William A. Powell, Frederick V. Hebard, John Scrivani, Caragh Heverly, Martin Cipollini, Brian Clark, Eric Evans, Bruce Levine, John E. Carlson, David Goodstein, Jane Grimwood, Jeremy Schmutz, Jason A. Holliday, John T. Lovell

## Abstract

Over a century after two introduced pathogens decimated American chestnut populations, breeding programs continue to incorporate resistance from Chinese chestnut to recover self-sustaining populations. Due to complex genetics of chestnut blight resistance, it is challenging to obtain trees with sufficient resistance and competitive growth. We developed high quality reference genomes for Chinese and American chestnut and leveraged large disease phenotype and genotype datasets to develop accurate genomic selection. Inoculation and simulation results indicate that resistance may be substantially increased in trees that inherited 70% to 100% of their genome from American chestnut. To facilitate gene editing, we integrated multiple lines of evidence to discover candidate alleles for blight resistance and susceptibility. These genomic resources provide a strong foundation to accelerate restoration of this iconic tree.

## Background

The demise of the American chestnut (*Castanea dentata*, (Marsh.) Borkh.) is a quintessential example of population collapse following introduction of non-native pathogens (*1*). The necrotrophic fungal pathogen causing chestnut blight (*Cryphonectria parasitica*, (Murr.) Barr) was introduced to North America on imported Asian chestnuts in the late 1800’s and early 1900’s (*2*, *3*). By 1950, chestnut blight killed billions of *C. dentata* stems from Maine to Mississippi (Fig. 1A), eliminating this once dominant tree from the forest canopy (*4*). While hundreds of millions of American chestnuts persist as root collar sprouts and understory seedlings (*5*), they rarely reproduce before succumbing to chestnut blight (Fig. 1B). As land development and environmental stressors continue to kill resprouting genotypes, the remaining population is a dwindling resource for restoration (*6*, *7*).

**Fig. 1.**
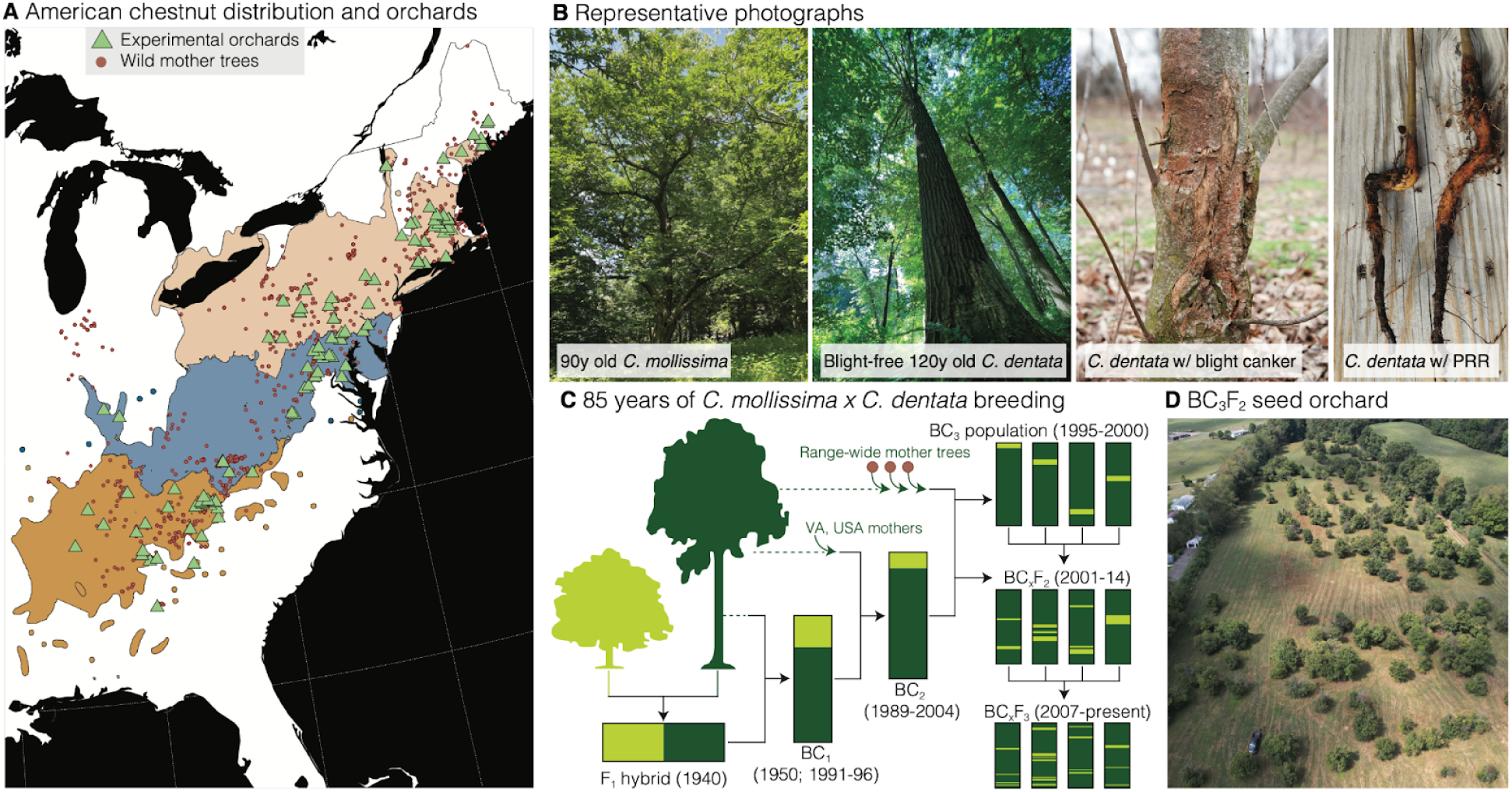
Biogeography of American chestnut and breeding efforts to improve tolerance to its non-native fungal pathogens. **(A)** The native distribution of *C. dentata* spans much of eastern temperate North America with the center of diversity residing along the Appalachian Mountains. Conservation efforts have involved breeding with rare reproductive *C. dentata* individuals (red dots) and growing progeny from these trees in orchards (green triangles) that span *C. dentata*’s three main climate regions (orange, blue and pink polygons). **(B)** Photographs (from left to right) of the ‘Mahogany’ *C. mollissima* tree at the Connecticut Agricultural Experiment Station; a *C. dentata* individual in the Arboretum of Tervuren in Belgium (Photo: A. Sproul-Lattimer); a ‘sunken’ blight canker on a 4 year old *C. dentata* stem; roots of two seedlings with significant Phytophthora root rot. **(C)** A schematic of the TACF hybrid/backcross breeding program; *C. dentata* (dark green) and *C. mollissima* (light green) proportional contributions are shown in the F_1_ and first two BC generations. Cartoons in the BC_3_F_3_ show what a single chromosome of segregants may look like. All wild mother trees (panel A) are parents of the BC_2_/BC_3_ population. **(D)** An overhead image of a representative orchard in Meadowview, VA taken by an unmanned aerial vehicle. The orchard was originally evenly planted - the sparsity is due to culling of trees with inadequate blight resistance.

Chestnut blight is not the only invasive pathogen threatening *C. dentata*. *Phytophthora cinnamomi* (Rands), an oomycete that causes Phytophthora root rot (PRR), was introduced into North America in the 1820s and killed many *C. dentata* and related Allegheny chinquapins (*Castanea pumila*, (L.) Mill.) (*8*, *9*). Although cold winter temperatures historically limited its impacts to the Southeastern U.S. < 40°N, PRR is expected to be more of a threat to *C. dentata* across its entire historical range under warming climates (*10*).

There has been over a century of effort to generate *C. dentata* varieties resistant to chestnut blight and PRR (*11*). Reforestation with disease-resistant *C. dentata* varieties has many potential benefits including the return of a regular mast of chestnuts for wildlife and human consumption (*12*), restoration of Appalachian strip mine sites (*13*), and carbon sequestration (*14*). In addition to disease resistance, restoration populations need to have the canopy competitiveness of pre-blight populations (*15*) and sufficient genetic diversity to adapt to a wide geographic range and changing climates (*7*).

### Potential pre-adapted sources of pathogen resistance

One source of resistance to chestnut blight is the handful of reproductively mature, large surviving American chestnuts (‘LSAs’) still growing in native forests. Starting in the 1980s the American Chestnut Foundation (TACF) and the American Chestnut Cooperators Foundation conducted controlled pollinations between these isolated LSAs (Fig. 1A-B), which are trees infected with blight for > 10 years with main stems > 25 cm in diameter (*16*). However, expectations from LSA-only breeding should be tempered: LSAs exhibit only moderate blight resistance, lack PRR resistance, and represent an extremely bottlenecked sample of genetic diversity (*16*, *17*).

Since Chinese chestnut (*C. mollissima*, Blume) and Japanese chestnut (*C. crenata*, Siebold & Zucc.) populations co-evolved with *C. parasitica* and *P. cinnamomi* and are resistant to these pathogens (*9*, *18*), interspecific hybridization efforts starting in the 1920s sought to imbue *C. dentata* hybrid populations with resistance alleles (Fig. 1C). Despite their strong pathogen resistance, many first generation (F_1_) hybrids lacked the maximum height growth of *C. dentata* (Fig. 1B) (*19*, *20*). Additional generations of backcrossing to *C. dentata* offered a means to recover dominant growth and was pursued by TACF under the hypothesis that a few major effect alleles from Chinese chestnut underlie blight and PRR resistance (*21*, *22*). While advanced backcross progeny are morphologically similar to American chestnut (*23*) and have enhanced blight and PRR resistance (*11*, *24*), molecular analysis revealed a tradeoff between blight resistance and *C. dentata* ancestry, implying that blight resistance is a quantitative trait (*25*). This conclusion is supported by genetic mapping experiments (Fig. 1D), which found quantitative trait loci (QTL) for blight resistance on all 12 chromosomes (*26*). Therefore, a key challenge of the American chestnut hybrid breeding program is to further improve pathogen resistance while simultaneously enhancing forest competitiveness (*11*, *25*).

Biotechnology offers a complementary restoration strategy to backcross breeding. Transgenic insertion of a wheat oxalate oxidase gene (OxO) into American chestnut has thus far shown the most promise for improving blight resistance (*27*). Federal regulatory review of transgenic trees expressing OxO is underway and these trees are currently being evaluated for blight resistance and growth in permitted field trials (*27*, *28*). Genome editing to knock out *C. dentata* susceptibility alleles or to insert multiple resistance alleles from Asian chestnut species may further enhance disease resistance, but will be challenging given the quantitative architecture of blight resistance and the long time frames (5-10 years) to validate candidate alleles in trees.

### Genomic foundations for American chestnut restoration

Breeding and biotechnology to improve disease resistance in American chestnuts were historically limited by a lack of genome resources. Here, we developed chromosome-scale *Castanea* genomes for three important founder trees: our *C. dentata* genome is from ‘Ellis-1’ (hereon “Ellis”), the tree that was transformed with the wheat OxO gene (*28*), and the two *C. mollissima* genomes are from the ‘Mahogany’ and ‘Nanking’ trees that contributed blight and PRR resistance to TACF’s breeding program (*11*) (Fig. 1B-C).

To construct the genomes, we assembled 112X - 125X PacBio sequencing, scaffolded the contigs into chromosomes with 76X - 79X HiC, and polished the assemblies with 50X-100X Illumina reads. Differences in sequencing technology necessitated a single haplotype build for ‘Ellis’, whereas we generated haplotype-resolved assemblies for ‘Mahogany’ and ‘Nanking’. The genome annotations encompass 25.2k-31.2k protein coding genes per assembly and were generated from homology, *ab initio* evidence, and tissue- and condition-specific RNA-seq support. The five assemblies are accurate (QV range = Q48-Q58), complete (BUSCO = 97.7-98.3%) and contiguous (contig N_50_ = 12-53Mb, Fig. 2A-B), indicating these are the highest quality *Castanea* genomes available (*29–34*).

**Fig. 2.**
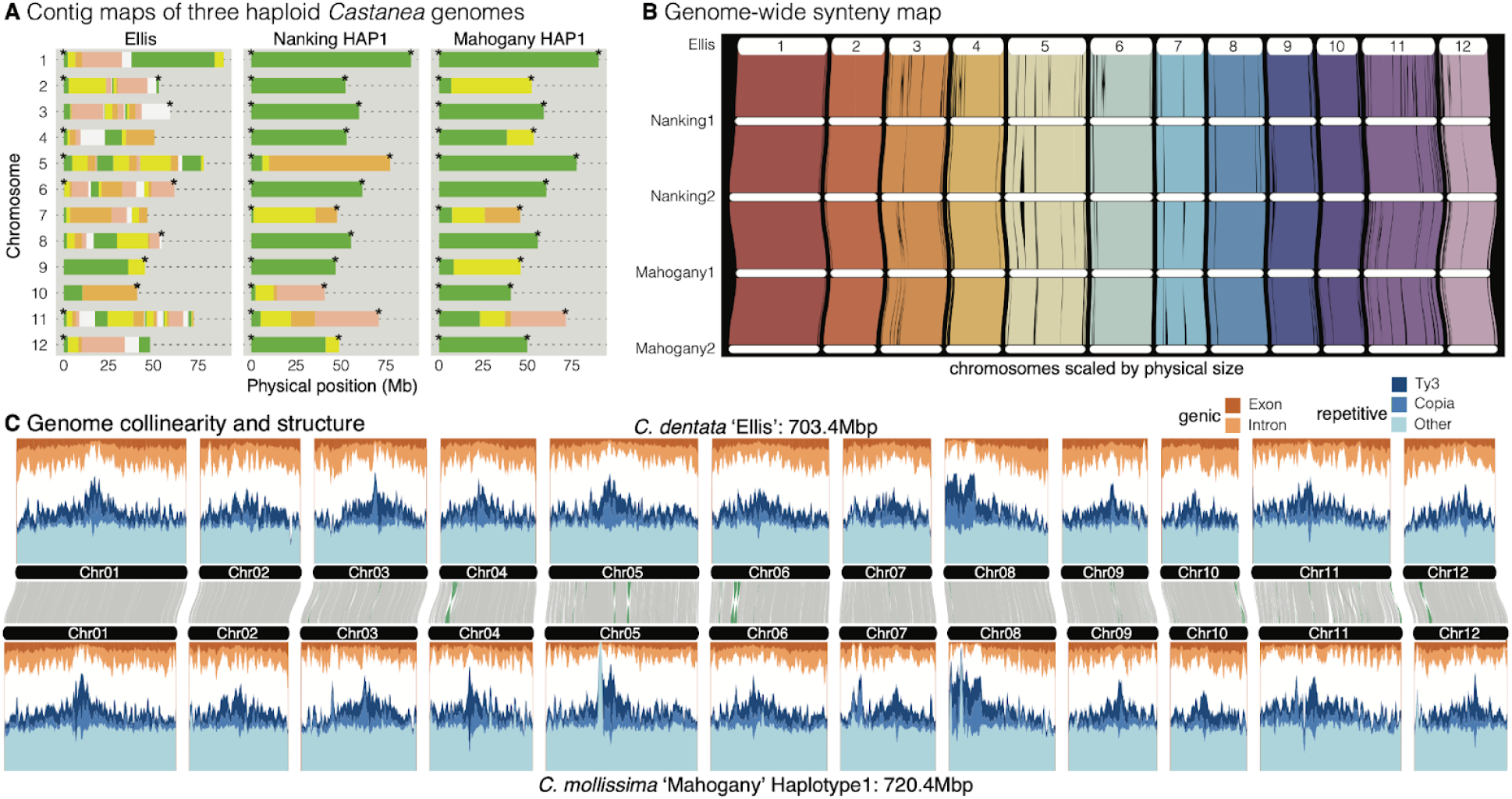
Genome structure and assembly for three *Castanea* genotypes. **(A)** Contig maps, as produced by GENESPACE showing the positions of contig breaks (gaps = where colors change) and telomeres (*) for three of the five genome assemblies. **(B)** DEEPSPACE riparian plot showing syntenic positions across all five assemblies. **(C)** Gene, repeat, and unannotated (white) sliding window densities in 1Mb 900-kb overlapping windows are connected by the syntenic links as were used to generate panel B; inverted syntenic blocks are highlighted in green.

Despite over 20 million years of divergence between *C. mollissima* and *C. dentata* (*35*), we expected these genomes to be highly conserved because of the high degree of synteny among the Fagaceae (*36*). As expected, there was strong sequence collinearity between species (mean = 662.0Mb, 94.7% of Ellis genome) (Fig. 2B-C). Genome structure within *C. mollissima* was even more conserved (mean = 3.50Mb of inverted sequence), and the two Nanking haplotypes only segregate for a single 110kb inversion. Furthermore, 90.7% (120,244) of genes are in families that span all five genomes. Only 6.9% (9,094) and 2.4% (3,140) of the gene models are in *C. dentata* and *C. mollissima* species-specific families, respectively. This similar genomic organization provides a strong foundation for comparative genomics and discovery of candidate resistance or susceptibility alleles.

### An expansive campaign to quantify pathogen resistance in American chestnut hybrid populations

In 1989, TACF began introgressing blight and PRR resistance via backcrossing the ‘Clapper’ BC_1_ ([*C. dentata* x *C. mollissima*] x *C. dentata*) and ‘Graves’ F_1_ (*C. dentata* x *C. mollissima*) founders with 61 flowering *C. dentata* trees from southwest Virginia (Fig. 1C) (*11*). These populations were subsequently diversified in a citizen-science led effort to breed selected backcross progeny with 344 additional flowering *C. dentata* trees ranging from coastal Maine to northern Alabama. Selected BC_2_ and BC_3_ progeny were intercrossed to generate large BC_x_F_2_ populations that segregate blight and PRR resistance (Fig. 1C-D). In 35 years of breeding, 49 additional *C. mollissima*, 16 LSA, and seven *C. crenata* sources of blight and PRR resistance were incorporated via backcrossing to 144 additional *C. dentata* parents.

To build a foundation for continued resistance gains, we phenotyped 5,501 trees aged 5-96 years (mean = 15 years), for eight stem traits indicative of blight susceptibility at 91 orchards across the eastern USA (Fig 1A). We summed blight resistance trait values and scaled the additive genetic variation in this ‘index’ from 0 (equal to average of susceptible *C. dentata* controls) to 100 (mean of resistant *C. mollissima*). In addition to simplifying interpretation, the blight resistance index had higher heritability (*h^2^* = 0.42 ± 0.03 s.e.) than the individual resistance variables (*h^2^* = 0.22 ± 0.05 to 0.31 ± 0.04 s.e.; Fig. S1). To assay PRR resistance, we inoculated first year seedling progeny with *P. cinnamomi* in contained trials from 2005 to 2023 and generated a 0 (*C. dentata* mean) to 100 (*C. mollissima* mean) scaled PRR survival index. Family mean PRR survival heritability was very high (median *n* per family = 28, *h^2^* = 0.97 ± 0.01 s.e.) among 26,288 open pollinated progeny from 662 mothers (Fig. S1) (*24*). Significant heritable variation for both blight and PRR resistance provides the basis for continued gains through selective breeding.

### Genomic selection to balance gains in pathogen resistance with forest competitiveness

Despite heritable resistance, phenotypic selection is slow in chestnut trees, requiring > 5 years to obtain reliable estimates of blight resistance (*37*). To accelerate gains, we implemented genomic selection, a technique that leverages genetic relationships to predict trait values (*38*), enabling selection of the most resistant seedlings prior to planting in seed orchards (*39*). As a first step towards building accurate genomic selection models, we mapped genotyping-by-sequencing (GBS) libraries of 5,003 trees to our ‘Ellis’ reference. From the resulting 93,333 single nucleotide polymorphisms (SNPs), we predicted the spectrum of Chinese and American chestnut ancestry across 12 breeding generations (Fig. 3A; Fig. S2). Furthermore, we estimated additive genetic variation (breeding values) for blight resistance (*n* = 6,585) and PRR survival (*n* = 2,208) indices across individuals that were phenotyped and/or genotyped.

**Fig. 3.**
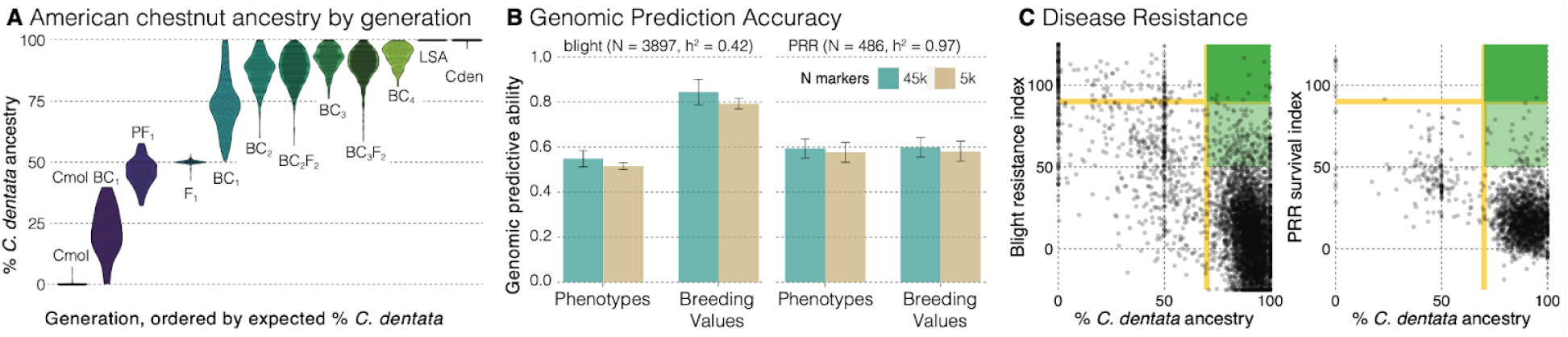
Hybrid species ancestry and genomic prediction of resistance to chestnut blight and PRR. **(A)** *C. dentata* ancestry was estimated from molecular markers for 12 breeding generations, including parental species (Cmol = *C. mollissima* and Cden = *C. dentata*), LSA (large surviving *C. dentata*), F_1_ (*C. dentata* x *C. mollissima*), PF_1_ (Pseudo-F_1_ = *C. mollissima* x BC_x_), and backcross (BC_x_/BC_x_F_x_) progeny. **(B)** Genomic prediction accuracy to predict additive genetic variation (breeding values) and phenotypes for the blight resistance index and PRR survival rates. Differences in training population size and heritability for these traits may explain differences in genomic prediction accuracy. **(C)** Correlations between individual tree measures of *C. dentata* ancestry and scaled blight/PRR (0 = mean of *C. dentata*; 100 = mean of C*. mollissima*); resistance values higher than 125 or lower than −25 are plotted at these axis limits to improve visualization. The dark green box represents the desired breeding outcome of plants with high *C. dentata* ancestry and high resistance. The light green boxes highlight trees with intermediate to high levels of blight resistance and majority *C. dentata* ancestry to select for breeding.

To assess genomic selection accuracy, we masked resistance phenotypes for five subsets of 20% of genotyped and phenotyped individuals. Our genomic selection models had significant prediction accuracy for both blight resistance (*r_phenotypes_* = 0.56 ± 0.03 s.d., *r_breeding values_* = 0.85 ± 0.05, *n* = 3,897) and PRR survival (*r_phenotypes_* = 0.60 ± 0.04, *r_breeding values_* = 0.65 ± 0.04, *n* = 486; Fig. 3B) indices with less than 1% bias in the prediction of phenotypes. Prediction with a genetic relationship matrix calculated from ∼5k markers incurred <10% loss in accuracy, indicating that genotyping costs may be reduced by the development of a lower density marker panel (*40*) (Fig. 3B).

While genomic selection models were highly predictive, improved pathogen resistance alone is not enough — chestnut trees must have competitive growth in native forests. As genetic control of growth is often complex in trees (*41*) and we cannot measure blight-free growth, we opted to use *C. dentata* ancestry proportions as a proxy for growth potential. We observed negative correlations between *C. dentata* ancestry and blight resistance (*r* = −0.65, *n* = 4,922, *P* < 0.0001) and PRR survival (*r* = −0.53, *n* = 2,051, *P* < 0.0001) indices (Fig. 3C). In a closed canopy stand containing LSA, backcross, and *C. mollissima* individuals (Lesesne State Forest, VA), height growth of the 30% fastest growing backcross trees was intermediate between that of LSA and *C. mollissima* individuals (Fig. S3), indicating that additional selection is required to fully recover competitive growth in backcross populations.

To further dissect the blight resistance-*C. dentata* ancestry tradeoff, we inoculated 1,015 seedling progeny from 30 families and observed a strong tradeoff between progeny mean blight resistance and the average *C. dentata* ancestry of the parents (r = −0.83; Fig. S4; Table S1). However, four *C. dentata* backcross families had mean blight resistance ratings ranging from 56 to 85 and majority parental *C. dentata* ancestry ranging from 64% to 68%. This indicates that medium to high levels of blight resistance are obtainable in hybrids with ∼70% *C. dentata* ancestry.

### The genetic architecture of pathogen resistance in American chestnut hybrid populations

The tradeoff between *C. dentata* ancestry and disease resistance implies that it may be difficult to breed trees with enough resistance for long term survival while retaining competitive growth. If resistance had an infinitesimal genetic architecture, variance explained per chromosome would be strongly correlated with chromosome size (*42*), and individual QTLs would not significantly explain resistance beyond simple *C. mollissima* ancestry proportions. For the blight resistance index, six of 12 chromosomes explained significant proportions of the variance (Table S2) and the correlation between chromosome length and variance explained was not significant (Spearman’s ρ = 0.30, *P* = 0.34; Fig. S5). For PRR survival, only one chromosome (Chr05) explained a significant proportion of the variance in survival (h^2^_chr_ = 0.83 ± 0.06; Fig. S5; Table S2). These results support a complex, but not infinitesimal genetic architecture for blight resistance and potentially a major gene architecture for PRR resistance.

To identify the largest effect loci, we conducted QTL mapping by associating linkage blocks of *C. mollissima* and *C. dentata* ancestry to resistance variation. For blight resistance, seven loci were detected on chromosomes 3, 5, 10, and 12 in the two largest backcross populations (*n*_Clapper_ = 1,475, *n*_Graves_ = 1,275; Fig. 4). Inheriting a *C. mollissima* allele at these loci was estimated to increase blight resistance index values by 8 to 11 units. For PRR resistance, we detected two large effect loci on Chr05, where inheriting a *C. mollissima* allele was estimated to improve the PRR survival index by 19 and 24 units (Fig. 4). The current population frequency of *C. mollissima* alleles among QTLs varied between 8% and 21% indicating there is opportunity to improve resistance via selection. While no overlap was detected in blight resistance QTLs among mapping populations (Fig. S6), there was a higher than expected frequency of *C. mollissima* alleles across QTLs on chromosomes 7, 10 and 12 among trees with blight resistance >50 from both the ‘Clapper’ and ‘Graves’ populations (Fig. 4; Fig. S7). This implies that there are likely to be both common and unique blight resistance loci across populations and breeding to stack resistance alleles may provide a path to enhanced resistance.

**Fig. 4.**
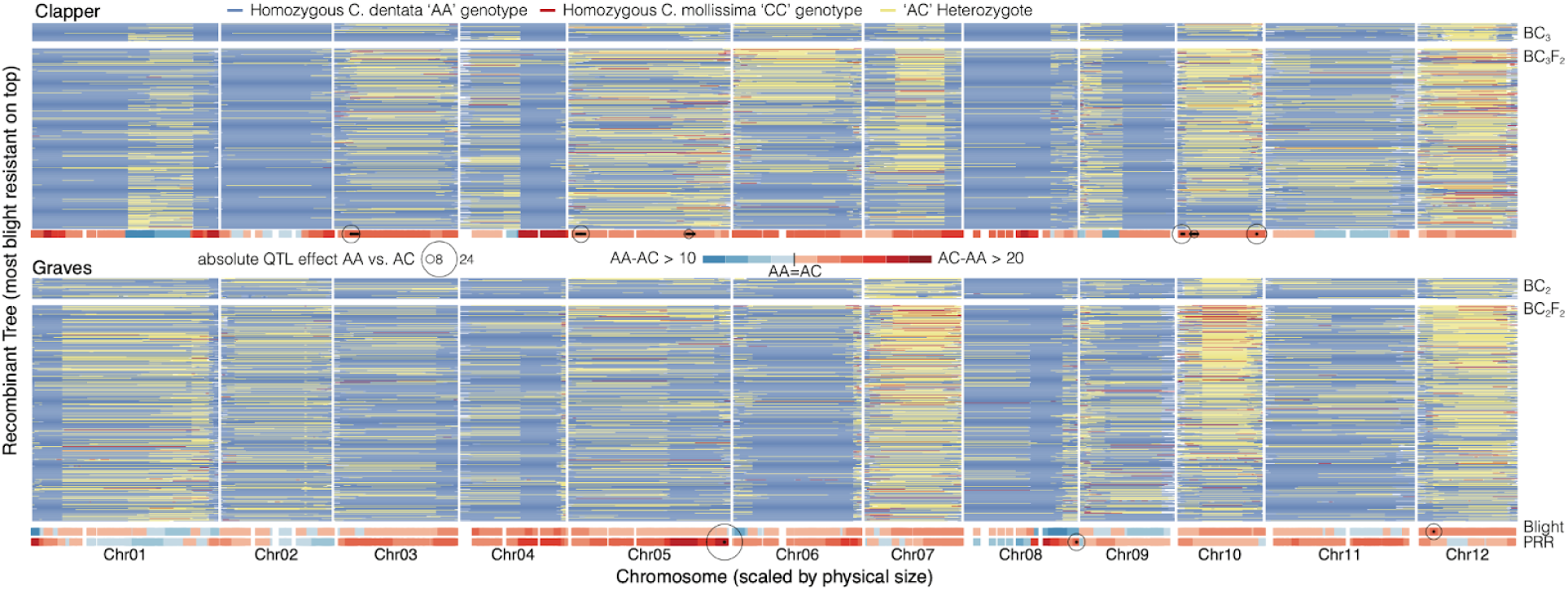
Genetic architecture of blight and PRR resistance in American chestnut hybrid backcross populations. Genotypic contributions of the progenitor species (red/yellow/blue map) are shown for 836 of the 2,750 recombinant progeny in the BC_2_/BC_3_ (smaller top facets) and BC_2_F_2_ and BC_3_F_2_ (larger bottom facets) for the two main mapping populations derived from the ‘Clapper’ and ‘Graves’ sources of resistance. We performed QTL mapping in each BC_x_F_x_ population. The red-blue horizontal bars below the genotype maps indicate ancestry effects on blight resistance and PRR survival indices. Red denotes regions where inheriting at least one allele from *C. mollissima* (‘AC’) enhances resistance, whereas blue denotes regions where inheriting both alleles from *C. dentata* (‘AA’) improves resistance. The circles indicate the point estimate and effect size of the QTL peak, and the horizontal black lines represent the QTL peak confidence interval. Allelic substitution effects (bottom horizontal bands) are estimated in units of the blight resistance and PRR survival indices.

### Next steps to improve pathogen resistance in American chestnut hybrid populations

‘Recurrent genomic selection’ (RGS) is one strategy to increase disease resistance while also selecting to maintain majority genome-wide *C. dentata* ancestry. Through multiple generations of recombination, RGS is expected to increase the frequency of resistance alleles (primarily from *C. mollissima*) in resistance-associated genome regions while maximizing *C. dentata* ancestry across the rest of the genome.

We evaluated a RGS strategy where parents with > 70% *C. dentata* ancestry on average and maximal resistance are intercrossed and the progeny are selected for enhanced resistance (Fig. 5A). To predict the efficacy of this strategy, we simulated blight and PRR resistance gains across four breeding tracks (Fig. 5B). The largest gains in blight resistance were predicted for intercrosses designed to maximize blight resistance (*track 1*). For the 20% selected progeny, blight resistance was predicted to vary between 81 and 108 (mean 86), and *C. dentata* ancestry varied between 70% and 85%. Balancing gains in blight and PRR resistance (*track* 2) resulted in lower gains for blight resistance (mean index 73) and PRR survival (mean index 60) compared to maximizing resistance to each disease individually (Fig 5B).

**Fig. 5.**
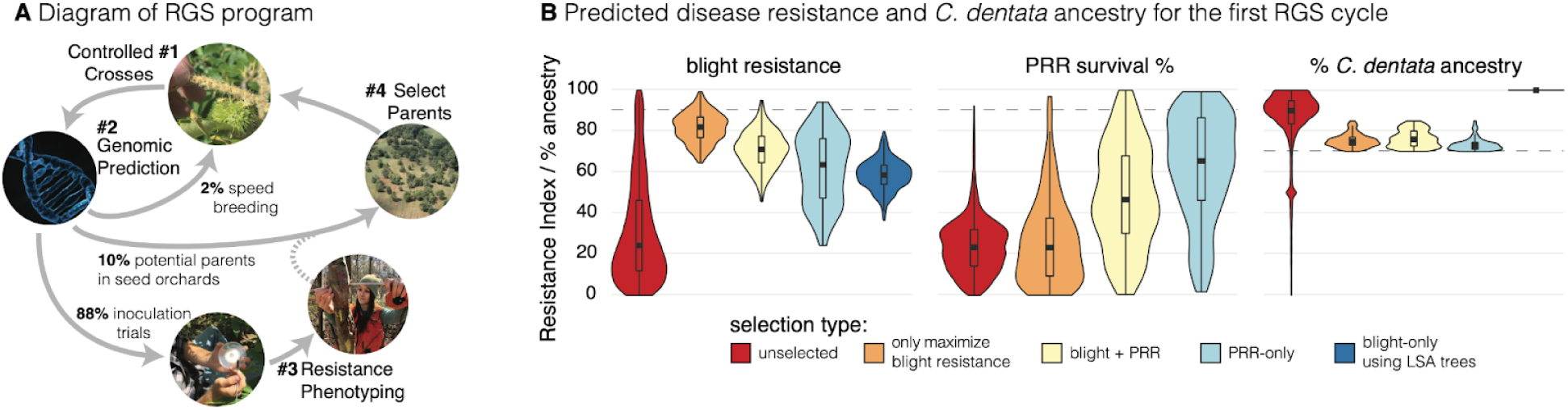
Next steps for breeding to improve blight and PRR resistance in American chestnut hybrid backcross populations. **(A)** Diagram of a single recurrent genomic selection (RGS) cycle. Percentages of individuals passed to each step in the cycle are indicated. The dashed line indicates that resistance phenotyping from relatives will be used to determine which parents should be used in controlled crosses. **(B)** Predicted genetic variation in the blight resistance index, PRR survival index, and American chestnut ancestry (%) over one cycle of controlled crosses and genomic selection. Breeding simulations were segmented into four tracks to 1) maximize blight resistance, 2) balance blight and PRR resistance, and 3) maximize PRR resistance while selecting for basal levels of blight resistance and 4) maximize blight resistance in large surviving American chestnut populations.

Additional generations of selection may be required to improve resistance to both diseases. Selected parents typically begin producing seeds within 7-15 years in orchards (*43*). Breeding cycles may be shortened to 1 to 2 years with supplemental lighting and fertilizer treatments (*44*) (Fig 5A). However, such ‘speed breeding’ approaches are unlikely to generate large numbers of progeny for reforestation or to realize substantial gains from selection; hence, our main strategy will be planting and evaluating trees in orchards.

### Discovering candidate genes involved in resistance to *C. parasitica*

Gene editing has the potential to complement RGS to enhance blight resistance, but requires knowledge of the specific (mal)adaptive alleles that impact host responses to *C. parasitica*. Candidates are often identified through coding, copy number, or regulatory variation, proximity to QTL peaks, and similarity to resistance genes (*45*). Among 2,538 and 1,997 genes from the ‘Ellis’ and ‘Mahogany’ Hap1 genomes that map to blight resistance QTL intervals, 201 had significant interspecific differential expression responses to *C. parasitica*, 938 orthogroups had putative copy number variants, 1347 single copy orthologs had non-synonymous variants, and 173 *C. dentata* (100 *C. mollissima*) genes were annotated as resistance genes. These lists of candidates are far too long for functional validation.

We narrowed the list of candidates via two *a priori* hypotheses. First, we hypothesized that genetic variation in promoter sequences increases the expression of resistance alleles uniquely in *C. mollissima* relative to *C. dentata* (*46*). Thirty-seven genes had *C. mollissima* allele-specific upregulated expression in *C. mollissima* x *C. dentata* F_1_ hybrids at 3 or 10 days post inoculation (Table S3). Four of these genes were within blight QTL intervals including a fungal chitinase that has a plausible role in resistance (*47*). Second, necrotrophic fungal pathogens are known to secrete effector proteins which interact with ‘resistance’ genes in susceptible hosts to hijack the programmed cell death response and promote disease (*48*). As candidates for susceptibility, we identified 26 resistance genes across four QTL intervals from a Mahogany F_2_ mapping population that were present in the ‘Ellis’ and absent in both haplotypes of ‘Mahogany’ (Table S4). To aid in future identification of host targets of effectors, we identified 22 effectors from *C. parasitica* that were upregulated in *C. dentata* hosts and primarily targeted to the apoplast (Table S5). While not currently feasible at scale, it will soon be possible to knock out susceptibility alleles or insert resistance alleles (*49*, *50*), enabling introgression into diverse *C. dentata* germplasm.

### Conclusions

Genetic rescue of American chestnuts with continued threats from two invasive pathogens will be assisted by the integration of genome-informed breeding and biotechnology. Here, we demonstrate that generating hybrids with > 70% American chestnut ancestry and sufficient blight and PRR resistance is possible. Our reference genomes and the development of a high-throughput genotyping platform will provide the basis for recurrent genomic selection to improve disease resistance while maximizing *C. dentata* ancestry. As trees begin to exceed resistance thresholds for long-term survival, selection to increase maximum height growth form will become an important additional breeding objective. Biotechnological approaches to insert resistance alleles from *C. mollissima* or knock out susceptibility alleles in *C. dentata* are complementary strategies to improve blight resistance without sacrificing *C. dentata* ancestry. Developing efficient and multiplexed methods are critical next steps for candidate gene validation.

## ACKNOWLEDGEMENTS

Laura Barth and Dan Mckinnon collected tissue for the Chinese chestnut reference genome sequencing. Shanmugan Rajasekar and David Kundra performed high molecular weight DNA extractions for the reference genomes. TACF volunteers and interns Israel Golden, Mira Polishook, Vinny Varsalona, Lizzy Dvorak, Daniel Ott, Mike Aucott, and Jay Brenneman assisted with phenotyping. Analyses were performed on Virginia Tech Advanced Research Computing clusters. Copy editing was performed by Christine Wiese.

## Funding

Members and Donors of The American Chestnut Foundation (JW)

Oak Hill Fund (JW, JH)

Foundation for the Carolinas (JW, JL, JS, JG)

Colcom Foundation (JW, JS, JG)

U.S. Department of Agriculture National Institutes of Food and Agriculture projects 1018599, 1027966, and 1025004 (JH, JW)

U.S. Forest Service (JW, SF, VL)

Tucker Foundation (JW)

Ballyshannon Foundation (SF, JW)

Orentreich Foundation for the Advancement of Science (SF, JW)

Office of Science of the U.S. Department of Energy Contract No. DE-AC02-05CH1123 (JS, JG, JL)

## Author contributions

Conceptualization: JW, JL, JH

Methodology: JW, JM, VL, SF, QZ, JL, TE, BL, TS, TE, AS, JHC, SK, CP, AS, XX

Investigation: JW, JM, QZ, VL, SFF, JVC, KC, SH, CS, LG, EJ, TS, BJ, LW, LK, CW, PZ, SK, TZ, XX, FC, JS, JO, SJ, KM, ES, MW, LB, JJ, PS, AN, EC, FH, JS, CH, MC, BC, EE, BL, AT

Data Curation: JW, AS, JH, DG Visualization: JL, JW

Funding acquisition: JW, JH, SF, VL, KC

Project administration: JW, SF, VL, JH

Supervision: JW, JH, JL, SF, VL, AA, CDN, KC, JS, JG, JC, WP

Writing – original draft: JW, JL, JH

Writing – review & editing: JW, JL, JH, JM, FH, AA, BL, AT, SJ, JS, SK

## Competing interests

Authors declare that they have no competing interests.

## Data and materials availability

Data, results, and R scripts are deposited in Dryad https://doi.org/10.5061/dryad.4xgxd25mj. Raw genomic and RNA sequence data used to develop reference genomes can be accessed via NCBI bioproject PRJNA1147634. Chestnut reference genomes are accessible on Phytozome. Raw fastq from files from genotyping-by-sequencing can be accessed through NCBI under bioproject PRJNA507748.

## SUPPLEMENTARY MATERIALS

### MATERIALS AND METHODS

#### High molecular weight DNA extraction

High molecular weight (HMW) DNA for genome assembly was extracted from young leaves at Clemson University for *C. dentata* and Arizona Genomics Institute for *C. mollissima* genotypes. For the ‘Ellis1’ *C. dentata* genome, tissue culture-generated plantlets were grown at SUNY ESF in the dark to reduce accumulation of carbohydrates. For the ‘Mahogany’ and ‘Nanking’ *C. mollissima* genomes, newly emerging leaves from field-grown trees were wrapped in a shade cloth for 48 hours. Leaves were flash frozen in liquid N upon collection and stored at −80 C. The HMW DNA extraction was conducted by the protocol described in (*51*) with modifications.

Briefly, nuclei were isolated using Nuclear Isolation Buffer (NIB) composed of 0.5 M sucrose, 2% of polyvinylpyrrolidone (M.W.40,000), 0.5% Triton X-100, 10 mM KCl, 50 mM EDTA, 4 mM spermidine trihydrochloride, 1 mM spermine tetrahydrochloride in 10 mM Tris-HCl at pH 8.0. Before use, the NIB was supplemented with 0.1% of 2-Mercaptoethanol. The tissue weight to NIB buffer volume ratio was adjusted to 1:10. Pelleted nuclei were washed with NIB supplemented with 0.5% Triton X-100 and resuspended in CTAB lysis buffer (2% CTAB, 2M NaCl,100 mM EDTA, 100 mM Tris-HCl, pH 8.0) in the presence of Proteinase K (0.45 mg /ml) either overnight on ice or for 2 hours at 65C. DNA was phase-separated with equal volume of chloroform and precipitated with 0.7 volume of cold isopropanol. The DNA was washed in 70% ethanol, dried at room temperature, and resuspended in 0.01 M tris HCl, pH 8.0. Then, the DNA was treated with 3 ul of the Ambion® RNase Cocktail™ (Thermo Fisher Scientific), for 30 min at 37°C, followed by chloroform: isoamyl alcohol (24:1 vol/vol) extraction and precipitation with two volumes of ethanol. The DNA was resuspended in 100 ul of 0.01 M Tris-HCl, pH 8.0. The quality and integrity of the DNA was evaluated using a Qubit 2 Fluorometer (Thermo Fisher Scientific Inc., USA), a NanoDrop ND-8000 spectrophotometer (Thermo Fisher Scientific Inc., USA) followed by Pulse Field Gel Electrophoresis in agarose gel.

#### Genome assembly: *C. dentata* ‘Ellis’

We produced 118.58X coverage of PacBio continuous long reads (CLR), which were assembled with MECAT (*52*) and polished with Quiver (*53*). A total of 24 misjoins were corrected in the polished assembly using a 980-marker genetic map from the ‘Cranberry’ *C. dentata* tree (*54*).

Contigs were assembled into larger scaffolds with HiC data using the JUICER pipeline (*55*). Using the genetic map, a total of 160 joins were applied to the scaffolds. The final assembly contained 99.37% of the assembled sequence in 12 chromosomes. Chromosome assemblies were ordered and oriented with respect to the *Quercus robur* genome (http://www.oakgenome.fr/) and the telomeric sequence was found to be properly oriented in the assembly. Forty eight adjacent alternative haplotypes were collapsed using the longest common substring. The raw CLR data were used to correct 76,296 heterozygous SNPs and INDELs (2.8% of the 2,718,565 heterozygous phasing errors). A total of 533 homozygous SNPS and 137,372 homozygous INDELs were corrected across 662,414,933 callable bases using 100X Illumina reads (2×150, 500bp insert).

#### Genome assembly: *C. mollissima* ‘Mahogany’ and ‘Nanking

We produced 112.82-125.78X of single haplotype PacBio HiFi coverage where the average read length was 16,111-23,789 bp across the two genomes. Two haplotypes for each genome were assembled using HiFiAsm+HIC (*56*). The resulting sequence was polished using RACON (*57*). We searched for misjoins in the assemblies using HiC data, but none were identified. Contigs for both haplotypes were oriented, ordered, and joined with JUICER (*55*). Six joins were applied to form the final assembly in which the 12 chromosomes contained 99.27-99.66% of the assembled sequence. One adjacent alternative haplotype was collapsed in the assemblies using the longest common substring between the two haplotypes. A total of 966/1,655 heterozygous SNPs and INDELs were corrected using the HiFi data. Additionally, 298/554 homozygous SNPs and 12,027/13,402 homozygous INDELs were corrected using 50-51X Illumina reads (2×150, 400bp insert). No vectors or contaminants were found in the assembled sequence. Chromosomes were numbered and oriented using the V1 *C. dentata* release.

#### Genome annotations

To improve annotation of gene models, RNA sequencing was performed on female flowers, catkins, leaves, bark, and stems that were wounded and/or infected with *C. parasitica* using methods described below in ‘RNA extraction and sequencing.’ Gene models were annotated using our standard pipeline developed by the DOE Joint Genome Institute and Phytozome. Transcripts were assembled using PERTRAN (*58*) from 4.28 billion 2 × 150-bp stranded paired-end Illumina RNA-seq data. PERTRAN conducts genome-guided transcriptome short-read assembly via GSNAP (v2013-09-30) (*59*) and builds splice alignment graphs after alignment validation, realignment, and correction (*60*). Subsequently, PASA (v2.0.2) (*61*) was used to assemble 181,468, 148,147, 146,457, 153,886, and 153,743 transcripts for the ‘Ellis’, ‘Mahogany’ HAP1, ‘Mahogany’ HAP2, ‘Nanking’ HAP1, and ‘Nanking’ HAP2 genomes, respectively. The assembled transcripts were aligned to the repeat-soft-masked genomes with up to 2,000-bp extension on both ends unless extending into another locus on the same strand.

Repeats were identified with RepeatMasker v.4.1.3 (*62*), RepeatModeler (v.open1.0.11), and RepBase (*63*). Homology support was provided by alignments to seven publicly available genomes (*Arabidopsis thaliana*, soybean, sorghum, grape, peach, rice and *Setaria viridis*) and Swiss-Prot proteomes. Gene models were predicted by the homology-based predictors FGENESH+ (v.3.1.0) and FGENESH_EST (*64*), which uses expressed sequence tags to compute splice site and intron input instead of protein/translated open reading frames (ORFs). In addition, gene models were predicted using EXONERATE (v.2.4.0) (*65*) and AUGUSTUS (v.3.1.0) (*66*) trained with high confidence PASA assembly ORFs with intron hints from short read alignments. The selected gene models were subject to Pfam analysis and gene models with greater than 30% Pfam TE domains were removed. We also removed (1) incomplete, (2) low-homology-supported without full transcriptome support and (3) short single exon (<300 bp coding DNA sequences) without protein domain or transcript supported gene models.

#### Comparative genomics

We used GENESPACE v1.2.3 (*67*) and DEEPSPACE v0.0.1 (https://github.com/jtlovell/DEEPSPACE) in R v4.4.1 (*68*) to conduct all comparative genomics analyses. GENESPACE employs OrthoFinder v2.5.4 (*69*) to build orthogroups, then parses orthogroup-constrained protein-protein BLAST hits into syntenic blocks with MCScanX (*70*). This produced genome-wide synteny-constrained sets of orthologs across the five reference haplotypes. DEEPSPACE finds syntenic block coordinates by converting query genomes into windows and aligning these to the reference genomes with minimap2 v2.28 (*71*).

#### Blight phenotyping

A total of 5,501 trees from 91 orchard locations were phenotyped for 10 long term blight resistance and growth traits between 2018 and 2023. Over 80% of the trees were inoculated using the cork-borer, agar-disk method (*16*). Trees were inoculated with the weakly virulent SG2,3 strain and/or the strongly virulent EP155 strain of *Cryphonectria parasitica* when they were between 2 and 15 years old (average inoculation age = 8 years). Non-inoculated trees were phenotyped based on natural chestnut blight infections. Trees were phenotyped when they were an average age of 15 years for presence/absence traits indicative of blight susceptibility including a dead main stem (‘mainstemalive’), cankers > 15 cm in length (‘largecankers’), expanding cankers (‘blightcontained’), exposed wood (‘exposedwood’), *C. parasitica* fruiting bodies (‘sporulation’), and stump sprouts (‘stumpsprouts’). Prior to quantitative genetic analyses, the presence of susceptibility traits were coded as 0 and their absence was coded as 1 so trees lacking indicators of susceptibility had higher blight resistance estimates. Blight cankers were also rated based on whether they were sunken (0, a susceptibility response), swollen (1, a partially resistant response), or flat (2, indicative of higher resistance) (named ‘sunkenswollen’). The percent of the tree canopy that was dead as a result of blight infection was converted into canopy survival proportion (‘propcanopysurvival’). Tree heights in meters (‘height_m’) were measured with a clinometer (Suunto PM 5/360 PC Clinometer) or a laser hypsometer (Nikon Forestry Pro II Laser Rangefinder). The diameter of the largest living stem in centimeter (‘dbhlargest_cm’) was measured at breast height (1.37 meters).

#### Phytophthora root rot phenotyping

A total of 33,639 seedling progeny from 922 open pollinated *C. dentata* backcross families, resistant *C. mollissima*, and susceptible *C. dentata* controls were inoculated with *P. cinnamomi* in yearly trials from 2005 to 2023 (*24*). Seedlings were grown in soilless media (Fafard 3B mix) in D40 pots (Steuwe & Sons) in a completely randomized design in trials at North Carolina State University (Raleigh, NC) and the U.S. Forest Service Resistance Screening Center (Asheville, NC). For trials at Chestnut Return Farms (Seneca, SC), seedlings were planted in Rubbermaid 150-gallon tubs in a randomized complete block design. At age 3 to 4 months, seedlings were inoculated with *P. cinnamomi* isolates grown on sterilized vermiculite moistened with V8 broth (*72*) or on sterilized rice grains (*73*). Seedlings were inoculated with *P. cinnamomi* isolates from locations where the seedlings were to be planted to avoid moving isolates between locations. Seedling survival was recorded 16 to 24 weeks post inoculation. For all trials except those at North Carolina State University, roots were rated for root rot severity with the following system: 0 = no lesions on roots, plant healthy; 1 = minimal lesions limited to secondary roots; 2 = any lesions on the tap root or extensive lesions on secondary roots; 3 = severe root rot, plant dead (*74*). Descendants of the ‘Clapper’ BC_1_ tree had little to no PRR resistance (*24*); therefore, progeny from this source of resistance were omitted from subsequent quantitative genetic analyses. Analyses were performed on a subset of 662 families (27,464 progeny) where 10 to 521 individuals per family (median 28 individuals) were phenotyped. Analyzed backcross families descended from the ‘Graves’ F_1_ (458 families), *C. mollissima* ‘Mahogany’ (36 families), *C. mollissima* ‘Nanking’ (43 families), and other sources of resistance (22 families). Twenty two susceptible *C. dentata* families and 14 resistant *C. mollissima* families were included in yearly trials as controls.

#### Genotyping

Genotyping-by-sequencing (GBS) was performed on 5,003 *C. dentata* backcross trees and wild type species controls. Library preparation and bioinformatic methods were described in detail in (*25*). Briefly, young leaves were collected, homogenized with liquid nitrogen, and genomic DNA extracted with a DNeasy Plant Mini Kit (Qiagen Company, Hilden, Germany). Samples were digested with ApeKI and barcoded with unique P1 and common P2 adapters ligated with T4 ligase. Libraries were subsequently PCR amplified with 14 cycles. Libraries were assessed on 1% agarose gels, quantified with a Qubit fluorometer (Thermo Fisher Scientific, Waltham, USA), and randomly assigned to pools of 96. Pools were size selected on a BluePippen instrument (Sage Science, Beverly, USA) and sequenced on an Illumina NovaSeq 6000 S-Prime flow cell in 2×150bp format at the Duke University School of Medicine. Fastq files were demultiplexed with the process_radtags function in Stacks (*75*) and aligned to the *C. dentata* ‘Ellis’ reference genome with bwa mem (*76*). Aligned sequence was converted to BAM format, sorted, and indexed with Samtools (*77*). Variants were called with GATK v4.2.6.1 (*78*) HaplotypeCaller (*79*) by creating gVCFs per individual and chromosome. The gVCFs were merged in batches of 50 samples with GenomicsDBImport and GenotypeGVCFs functions in GATK. The 50 sample VCF files were merged with bcftools to create a dataset comprising all samples and chromosomes. The whole population VCF was filtered to remove sites that did not meet the following quality thresholds: MQ<40.00, QD<2.0, FS>40.000, MQRankSum< −12.500, ReadPosRankSum<-8.000, and SOR>3.0. Variants detected in repetitive regions of the ‘Ellis’ *C. dentata* genome were masked. Remaining variants were further filtered with bcftools to retain a total of 93,333 biallelic SNPs with phred quality scores > 30 and less than 20% missing data across individuals. The filtered VCF was imputed and phased in Beagle v 5.4 (*80*, *81*).

#### Species ancestry inference

*Castanea* species ancestry in hybrids was inferred in Ancestry_HMM (*82*) using whole genome sequence (WGS) data from 21 *C. dentata*, 23 *C. mollissima*, 15 *C. crenata*, 16 *C. sativa*, 27 *C. pumila*, 20 *C. henryi,* 17 *C. seguinii* obtained from (*83*) and (*33*). Sequence was aligned to the *C. dentata* ‘Ellis’ reference genome and variant calling was performed using the same methods as for GBS to generate a final VCF containing 143,933,272 variants. Biallelic SNPs genotyped with GBS (N SNPs = 93,333) were subset from the WGS data and these datasets were merged with bcftools. Separate Ancestry_HMM runs were performed for the following putative ancestry combinations: *C. dentata* v. *C. mollissima* (4,504 trees), *C. dentata* v. *C. mollissima* v. *C. crenata* (479 trees), *C. dentata* v. *C. sativa* v. *C. crenata* (6 trees), and *C. dentata* v. *C. pumila* v. *C. henryi* (14 trees). Prior to running the analyses, per locus Fst between reference populations was estimated with the R package ‘hierfstat’ (*84*). Ancestry informative markers with Fst > 0.7 between reference populations and > 1 Kb physical distance between markers were subset and reference and alternative alleles were counted for each reference population. Genetic distances between markers were estimated by building two parental genetic maps in R/QTL (*85*) from GBS data from 125 full sibling progeny of two wild-type *C. dentata* (GMBig x Horn) using the pseudo-test cross method (*86*). The ‘Horn’ map, composed of 1,003 markers and covering 99.5% of the genome length, was used to interpolate the genetic map positions for the ancestry informative markers using the program predictGMAP (https://github.com/szpiech/predictGMAP). For the *C. mollissima* v. *C. dentata* ancestry estimates, Ancestry_HMM was run by specifying average global ancestry proportions of 0.88 for *C. dentata* and 0.12 for *C. mollissima*, four pulses of *C. dentata* ancestry (corresponding to F_1_ through BC_3_ generations) and one pulse of 5% *C. mollissima* ancestry (corresponding to intercrossing in the BC_3_F_2_ generation). For the three way ancestry contrasts, population mean ancestry proportions were estimated first with ADMIXTURE (*87*) and three pulses of ancestry were specified corresponding to these proportions. Ancestry calls (*i.e.*, AA = homozygous American chestnut, AC = heterozygous, and CC = homozygous Chinese chestnut) were made if a genotype probability was greater than 0.5. Start and end coordinates for species ancestry blocks were obtained with the ‘add_rle’ function from GENESPACE (*67*) after converting potentially spurious short ancestry runs (< 20 markers) to missing. Global *C. dentata* ancestry percentages for individual trees were estimated as % dentata = (2ΣL_AA_ + ΣL_AC_)/2L *100, where ΣL_AA_ and ΣL_AC_ are the summed lengths of AA of the AC genotype blocks and L is the total length of the genome in Mb.

#### Pedigree reconstruction

Pedigree records for phenotyped or genotyped trees, their parents, and previous generations back to founders were obtained from The American Chestnut Foundation’s online database, dentataBase (https://acf.herokuapp.com/). Complete maternal and paternal pedigree records were available for 1,965 of 5,501 of the blight phenotyped trees that were progeny of controlled pollinations. Paternity was assigned for an additional 2,007 trees from the genotypes of 288 candidate fathers using the program FAPs v. 2.6.4 (*88*). A total of 182 SNPs with less than 10% missing genotype data, that were spaced at least 1.1 Mb apart, and had expected levels of heterozygosity were selected for paternity analysis. The FAPs ‘paternity_array’ and ‘sibship_cluster’ functions were run assuming a mutation rate of 0.0015 and a maximum of four paternity clashes. Paternity was assigned to the highest probability father for 1110 progeny and an additional 897 progeny that were missing a single highest probability father were assigned to their second highest probability father.

#### Heritability and breeding value estimation

The additive genetic component of resistance traits and growth was estimated in ASReml-R v. 4.2 (*89*) with the mixed model *y* = *Xb* + *Zu* + *e*. The vector of phenotypes (y) is a function of fixed effects (b), random additive genetic effects (*i.e.*, breeding values) u ∼ N(0, σ^2^_a_ **H**), and residual error e ∼ N(0, σ^2^_e_ **I**). X and Z are incidence matrices relating phenotypes to fixed and random effects, respectively.

Binary traits were analyzed with binomial models and all other traits were modeled as continuous gaussian traits. PRR survival and root lesion severity ratings were strongly correlated; therefore, a bivariate model was used to estimate breeding values for these traits. A multivariate model was not computationally feasible for the 10 blight resistance and growth variables and separate univariate models were used to estimate breeding values for these traits.

To account for environmental effects on blight and growth traits, inoculation status, age class, climate, and soil variables important for *C. dentata* habitat suitability (*90*) were modeled as fixed factors. Average monthly maximum temperature (‘tmax’) and average monthly precipitation (‘prec’) for each orchard were retrieved using the ‘raster’ R package (*91*). Soil pH and soil percent sand content were retrieved from XPolaris in R (*92*). To model continuous tree ages, soil, and climate variables as discrete fixed factors, values were assigned to quartiles based on the distribution across individuals and orchard locations. To account for fixed effects on PRR traits, pathogen isolate, site, and year were combined into a single factor. All fixed factors had significant effects on the phenotypes (Wald F-test, P < 0.05); however, adjustment for these effects yielded variables that were strongly correlated with the unadjusted phenotypes (blight resistance index r_y,yadj_ = 0.95, PRR family mean survival r_y,yadj_ = 0.98).

Breeding values were estimated for both genotyped and non-genotyped individuals using single step genomic best linear unbiased prediction (ssGBLUP) (*93–95*). Pedigree (**A**) and genomic relationships (**G**) were blended into a hybrid relationship matrix (**H**) after scaling mean diagonal and off diagonal elements of **G** to those of **A** using the method of (*96*). Breeding values for the blight resistance were estimated for a total population of 6,585 trees including 3,897 individuals that were phenotyped and genotyped, 1,585 individuals that were phenotyped only, and 1,084 individuals that were genotyped only. Breeding values for family means of PRR survival and lesion severity were estimated for 2,208 trees, including 486 genotyped mothers with progeny phenotypes, 176 non-genotyped mothers with progeny phenotypes, and 1,546 individuals with genotype but no progeny test data.

We used the genomic relationship matrix (**G**) to estimate heritability (h^2^). Genomic relationships estimate Mendelian segregation within families and provide unbiased estimates of h^2^ (*97*). Using functions from the R package ‘ASRgenomics’(*98*), **G** was estimated with the method of (*99*) after pruning the SNP matrix to 48,266 markers with minor allele frequencies > 0.05 and genotypic correlations between markers < 0.9. Heritability for blight resistance traits was estimated as h^2^ = σ^2^_a_ / σ^2^_a+_ σ^2^_e_, where σ^2^_a_ is the additive genetic variance and σ^2^_e_ the error variance. For binary traits, σ^2^_e_ was assumed to equal the variance of the standard logistic distribution (π^2^/3 = 3.29) (*100*). For PRR traits, family mean heritability was estimated as h^2^ = σ^2^_a_ / σ^2^_a +_ σ^2^_e_ /n, where n is the harmonic mean of the number of progeny evaluated per mother tree.

#### Construction of a blight resistance index

A ‘blight resistance index’ was created to simplify interpretation and facilitate downstream genomic prediction and QTL mapping. To construct the index, individual variables were corrected for fixed factors and the residuals were scaled from 0 to 1 and weighted by multiplying by each trait’s heritability. Missing trait values were imputed in ‘phenix’ (*101*) from correlations among traits and pedigree relationships among trees. Weighted residuals were summed from the variables ‘mainstemalive’, ‘largecankers’, ‘blightcontained’, ‘exposedwood’, ‘sporulation’, ‘sunkenswollen’, and ‘stumpsprouts’. We selected this combination of traits because it was most strongly correlated with average blight canker ratings of progeny (see section on ‘progeny validation of blight resistance’). Blight resistance index values (*i*) for individual trees were scaled to 0 = mean for susceptible *C. dentata* and 100 = mean for resistant *C. mollissima* using the formula *i* = (*i*’ − µ_*am*_/(µ_*c*ℎ_− µ_*am*_), where µ_*am*_and µ_*c*ℎ_are the average unscaled index values (*i*’) for 86 susceptible *C. dentata* chestnuts and 92 resistant *C. mollissima* inoculated with *C. parasitica*.

#### Accuracy and bias of genomic prediction

Five-fold cross validation was performed to evaluate accuracy and bias in genomic prediction. Phenotypes were masked for 20% of genotyped and phenotyped individuals (Blight N = 3,897, PRR N = 486). Genomic estimated breeding values (GEBVs) were predicted from H-matrix relationships with remaining phenotyped individuals including those that were not genotyped. This procedure was repeated five times until GEBVs were obtained for the entire training population. Means and standard deviations for prediction performance metrics were calculated across the five folds. Predictive ability was estimated as the Pearson correlation between GEBVs and phenotypes after accounting for fixed factors (r_*gebv*, *yadj*_). Genomic prediction accuracy - the theoretical correlation between GEBVs and true breeding values (r_*gebv*, *bv*_) - was estimated by dividing predictive ability by the square root of trait heritability (*102*). To minimize bias in genomic prediction, cross validation was performed using different scaling factors for **G**^-1^ and **A**^-1^ varying from 0 to 1 (*103*). We selected the combination of scaling factors where the mean slope (b) of the regression of GEBVs (x) on y_adj_ (y) across five replicates of cross validation was 1 ± 0.01 (< 1% bias).

#### Progeny validation of blight resistance

To determine whether parental blight resistance index estimates were predictive of progeny blight resistance, 1,015 seedling progeny from 30 families were inoculated with *C. parastica*. Control families included three open pollinated (OP) *C. mollissima* families (resistant control), one *C. dentata* x *C. mollissima* F_1_ family (intermediate resistance control), and one OP *C. dentata* family (susceptible control). Test families included 10 OP *C. dentata* backcross families, 8 controlled pollinated (CP) backcross-F_2_ families, 6 quasi-BC_1_ (F_1_ x BC) families, and 1 LSA x LSA family. Seedlings were in 1.5 gallon pots for five months prior to inoculation in outdoor nurseries at the University of Tennessee Chattanooga (∼70% of the seedlings within families) and Berry College in Georgia (30% of the seedlings). Seedlings were inoculated by cutting off the apical leader of the seedlings and placing an agar disk containing inoculum from the highly virulent EP155 strain of *C. parasitica*. Ninety days post inoculation, orange zone canker lengths were measured to the nearest mm and the presence or absence of fungal fruiting bodies was noted (*104*, *105*). To control for outliers in the canker length distribution, canker severity ratings, on a 1 to 5 scale, were derived by assigning trees to quartiles from the canker length distribution and adding one to the rating if fungal fruiting bodies were present. Family mean canker ratings were scaled from 0 (*C. dentata* mean) to 100 (*C. mollissima* mean) and correlations were estimated with average blight resistance index and *C. dentata* ancestry of the parents.

#### Heritability by chromosome

To determine whether blight resistance index and PRR family mean survival had an infinitesimal genetic architecture or were likely controlled by major effect loci, correlations were estimated between heritable variance explained per chromosome and chromosome lengths. Additive genetic variance components for individual chromosomes was estimated in ASReml-R by fitting 12 individual chromosome models each with two G-matrices: one for the focal chromosome (σ^2^_chr_) and another for the remainder of the genome (σ^2^_g’_) (*42*, *106*). Heritability per chromosome was estimated as h^2^_chr_ = σ^2^_chr_ /(σ^2^_chr_ + σ^2^_g’_+ σ^2^_e_). To determine whether individual chromosome heritability was significantly greater than zero, likelihood ratio tests were performed where the log likelihood of the full model containing the focal chromosome was compared to that of a reduced model where genetic variance was estimated for the remainder of the genome.

#### Quantitative trait locus mapping

Quantitative trait loci associated with the blight resistance index and PRR survival were mapped in *C. dentata* backcross hybrid populations by associating variation in these traits with *C. mollissima* versus *C. dentata* ancestry calls from Ancestry_HMM (*82*, *107*). Blight resistance index QTL were mapped separately for 1,475 BC_3_F_2_ descendants of the ‘Clapper’ BC_1_ tree and 1,275 BC_2_F_2_ descendants of the ‘Graves’ F_1_ tree. PRR survival rates from 410 open pollinated (primarily BC_2_F_2_) families that descended from ‘Graves’ were used to map QTL for PRR resistance. Ancestry calls were pruned to 641 loci with a minimum recombination frequency between markers of 0.001 using the ‘dropSimilarMarkers’ function from ‘qtlTools’ (*108*). There was a low frequency of homozygous *C. mollissima* ancestry calls (3.7% of all calls). Therefore, to maximize power to detect QTLs, CC calls were converted to ‘AC’ and QTL analysis was performed specifying a backcross population type. A genetic map for the ancestry calls was estimated with the ‘est.map’ R/QTL function and Kosambi distances, assuming chromosomes were linkage groups and markers were ordered by physical position in the ‘Ellis’ *C. dentata* genome. The ‘scan1’ function from the R package ‘qtl2’ (*109*) was used to estimate LOD scores for marker-trait associations after controlling for polygenic kinship effects with the leave one chromosome out method (*110*). Significance thresholds for QTLs were estimated with the ‘scan1perm’ function and 1,000 permutations. Peak and border markers for QTL intervals were obtained with the ‘find_peaks’ and ‘lod_int’ functions specifying a peakdrop of one LOD unit. Genetic variance in resistance indices explained by the QTL intervals was estimated with R^2^ = 1 - 10^-2/N^ * ^LOD^, where N is the population size and LOD is the score for the peak marker (*111*). The effect of inheriting a *C. mollissima* allele at the QTL, in resistance index units, was estimated with the ‘scan1coef’ function. The genomic extent of the blight and PRR resistance QTL intervals mapped in this study were compared those from previous studies in American chestnut backcross populations (*26*, *54*) by blasting flanking sequences from peak and border markers against ‘Ellis’ *C .dentata* and both haplotypes from the ‘Mahogany’ and ‘Nanking’ *C. mollissima* genomes in Phytozome.

#### Simulated gains from genomic selection

Gains in the blight resistance index and PRR survival rates were simulated for one generation of controlled crosses between selected parents and genomic selection of the top 20% most resistant progeny. Three *C. dentata* backcross hybrid breeding tracks were simulated where biparental combinations with > 70% average *C. dentata* ancestry were selected to 1. maximize blight resistance (*i.e.,* 30 crosses where midparent blight resistance index was > 90), 2. balance gains in blight and PRR resistance (*i.e.,* 20 crosses where midparent blight resistance index > 60 & midparent PRR root lesion severity rating > 35), or maximize PRR resistance while maintaining a baseline level of blight resistance (*i.e.,* 10 crosses where midparent blight resistance index > 40 & midparent PRR root lesion severity rating > 45). A fourth breeding track maximized blight resistance among progeny of large surviving American chestnuts (*i.e.*, 10 crosses where midparent blight resistance index > 60 & both parents had 100% *C. dentata* ancestry). Simulated crosses were performed among selected parents from the central breeding region (*e.g.*, Virginia, Maryland, and Kentucky) (*7*). To minimize potential inbreeding, crosses were selected where coancestry coefficients between parents did not exceed 0.0625 and each parent was used in no more than three crosses. Genotypes of 100 progeny per cross were simulated with the ‘makeCross’ function in AlphaSimR (*112*) using phased parental genotypes from ∼93,333 biallelic SNPs and their interpolated genetic map positions. Genomewide *C. dentata* v. *C. mollissima* ancestry proportions of the simulated progeny were estimated with Ancestry_HMM (*82*, *107*). Genomic estimated breeding values (GEBVs) for the blight resistance index and family means for PRR root lesion severity ratings were predicted for the simulated progeny with ssGBLUP in ASReml-R v. 4.2. Training populations consisted of 3,897 genotyped trees phenotyped for blight resistance and 486 genotyped mother trees with PRR progeny test data. The GEBVs were scaled from 0 (*C. dentata* mean) to 100 (*C. mollissima* mean) as described previously. For breeding track 2 that balanced blight and PRR resistance a blight + PRR resistance index was created from the average breeding values of the blight resistance index and the scaled PRR root lesion severity rating. The top 20% progeny with > 70% *C. dentata* ancestry and the highest GEBVs for blight resistance (tracks 1 & 4), blight + PRR resistance (track 2), or PRR resistance (track 3) were selected. To predict realized gains from selection, true breeding values were simulated for the selected progeny by multiplying the GEBVs by genomic prediction accuracies estimated with cross-validation which were r_gebv,bv_ = 0.85 for blight resistance and r_gebv,bv_ = 0.65 for PRR resistance.

#### Experimental design for *C. parasitica* RNAseq timecourses

Species- and allele-specific responses to *C. parasitica* inoculation were compared at 3 and 10 days post inoculation. RNA from six biological replicates from *C. mollissima*, *C. dentata*, and F_1_ hybrids of these species were sequenced at each timepoint. Two genotypes per species or hybrid were sequenced and half of the biological replicates were derived from each genotype. The *C. dentata* genotypes included ‘Ellis’ and full sib progeny from trees from Tennessee (SM156 x TNCAN1). ‘Mahogany’ and ‘Nanking’ were the *C. mollissima* genotypes. The two F_1_ genotypes included KY115 and WWC70, which are progeny of two *C. dentata* trees from Virginia crossed with ‘Mahogany’ and ‘Nanking’ *C. mollissima* trees, respectively. To generate biological replicates, all genotypes were clonally propagated by whip and tongue grafting except the SM156 x TNCAN1 *C. dentata* progeny, where full siblings were used as biological replicates. Grafts and seedlings were grown in an outdoor nursery in two gallon pots at University of Tennessee Chattanooga for six months prior to inoculation. For each species and for F_1_ hybrids, transcript abundance was compared between two treatments: 1. stem wounding & inoculation with the EP155 strain of *C. parasitica* and 2. stem wounding without inoculation (control treatment). All trees were wounded by making a 1 cm long x 2 mm wide excision with a scalpel. Stem segments were sampled by cutting 0.5 cm above and below the wound and with Felco pruners and were immediately flash frozen in liquid nitrogen.

#### RNA extraction and sequencing

Total RNA was extracted at the Connecticut Agricultural Experiment Station after homogenizing tissue with a mortar and pestle in liquid nitrogen. Each RNA sample was extracted from 100-200 mg of frozen tissue in RNase free 5 ml tubes using a CTAB-based RNA extraction buffer using the protocol of (*113*) with the following modifications: Each sample was extracted with 3 ml of CTAB buffer with 120 µl of 20% SDS, extracted twice with chloroform:IAA, and LiCl precipitation was not performed. Samples that had at least 400 ng/µl nucleic acids based on Nanodrop 2000 (Thermo Scientific) readings were purified with an RNA extraction kit (PureLink RNA Mini Kit, Invitrogen, cat. No. 12183018A) combined with on-column DNase treatment (PureLink, Invitrogen, cat. 12185-010). RNA quality and quantity was estimated with Agilent TapeStation RNA Screen Tapes. Samples with < 200 ng/µl of RNA were vacuum-concentrated and the concentration and quality of these samples was re-evaluated with Invitrogen Qubit 4 fluorometer, Qubit BR RNA assays, and Qubit RNA IQ assays. Samples that had RNA Integrity Number (RIN) or RNA IQ score > 7.0, no DNA contamination, and > 200 ng/µl of RNA were sequenced. Messenger RNA was isolated for sequencing at HudsonAlpha Institute for Biotechnology using a Sciclone NGS robotic liquid handling system (PerkinElmer) and Illumina’s TruSeq Stranded mRNA HT Poly(A) RNA sample preparation kit. One μg per sample of total RNA was used as starting material and eight cycles of PCR were used for library amplification. The prepared libraries were quantified using KAPA Biosystems’ next-generation sequencing library qPCR kit and run on a Roche LightCycler 480 real-time PCR instrument. Sequencing of the flowcell was performed on the Illumina NovaSeq sequencer using NovaSeq XP v.1 reagent kits and an S4 flowcell, following a 2×150 bp indexed run recipe.

#### Species and allele-specific gene expression responses to *C. parasitica*

To contrast *Castanea* species gene expression responses to *C. parasitica*, reads from *C. dentata* were aligned to the ‘Ellis’ reference and reads from *C. mollissima* samples were aligned to ‘Mahogany’ hap1 genome assemblies using STAR 2.7.10a (*114*) after filtering sequencing adaptor and low quality sequences in FASTP v0.23.2 (*115*). To quantify allele-specific expression, reads from the *C. dentata* x *C. mollissima* F_1_ hybrids were competitively mapped to a concatenation of the ‘Ellis’ and ‘Mahogany’ references. Read counts for alleles were quantified by summing the reads that uniquely mapped to one genome with reads that mapped to both orthologs using the ‘featureCount’ function in the ‘Subread’ aligner (*116*). Expression responses to *C. parasitica* inoculation were quantified within species or allele in F_1_ hybrids at each timepoint by contrasting read counts from the inoculation and wounding treatments in DESeq2 (*117*). Species- and allele-specific responses were compared by contrasting the change in expression after inoculation for 19,312 single copy orthologs between the ‘Ellis’ and ‘Mahogany’ hap1 genomes inferred in GENESPACE (*67*). Expression differences were deemed significant for genes with |log_2_ fold change| > 1 and Benjamini-Hochberg false discovery rate < 0.05.

#### Identification of candidate genes in chestnut blight QTL intervals

Candidate genes for chestnut blight resistance were identified across seven blight resistance QTL identified in the current study and five QTL previously identified in *C. mollissima* x *C. dentata* F_2_ population (*26*). We identified genes from the ‘Ellis’ and ‘Mahogany’ hap1 genomes by intersecting the genomic features format genome (gff) gene & exon annotations with QTL boundaries using the ‘foverlaps’ function in ‘data.table’ (*118*). The list of genes in QTL intervals was intersected with species- and allele-specific expression datasets to identify genes with significant interspecific differences in the response to *C. parasitica* inoculation. Copy number variants (CNVs) and presence absence variants (PAVs) between the *C. dentata* and *C. mollissima* genomes were identified by comparing numbers of genes within orthogroups identified GENESPACE (*67*). Single copy orthologs with non-synonymous coding variation were identified by comparing the protein coding sequences between the ‘Ellis’ and ‘Mahogany’ hap1 genomes with ‘Biostrings’ (*119*). The program RGAugury (*120*) was used to specifically identify PAV and CNVs of ‘resistance gene analogs’ (RGAs) across the genomes.

#### Fungal effector identification and *in planta* expression

The program EffectorP v. 3 (*121*) was used to identify 274 putative effectors from the *C. parastica* secretome. Prior to running EffectorP, secreted proteins from the were identified from version 2 of the *C. parasitica* genome (*122*) in SignalP v. 6 (*123*). To identify *C. parasitica* effectors that were upregulated in *C. mollissima* and *C. dentata* hosts, transcript abundance *in planta* was compared to two *in vitro* control samples where RNA was sequenced from the EP155 strain grown for a week in the dark at 24°C on potato dextrose agar and in potato dextrose broth. Genes from *C. parasitica* upregulated within each host at each timepoint were identified in DESeq2 (*117*) as those with log2fold change > 1 and Benjamini-Hochberg P < 0.05 in the *in planta* - *in vitro* contrasts.

## SUPPLEMENTARY FIGURES AND TABLES

**Fig. S1.**
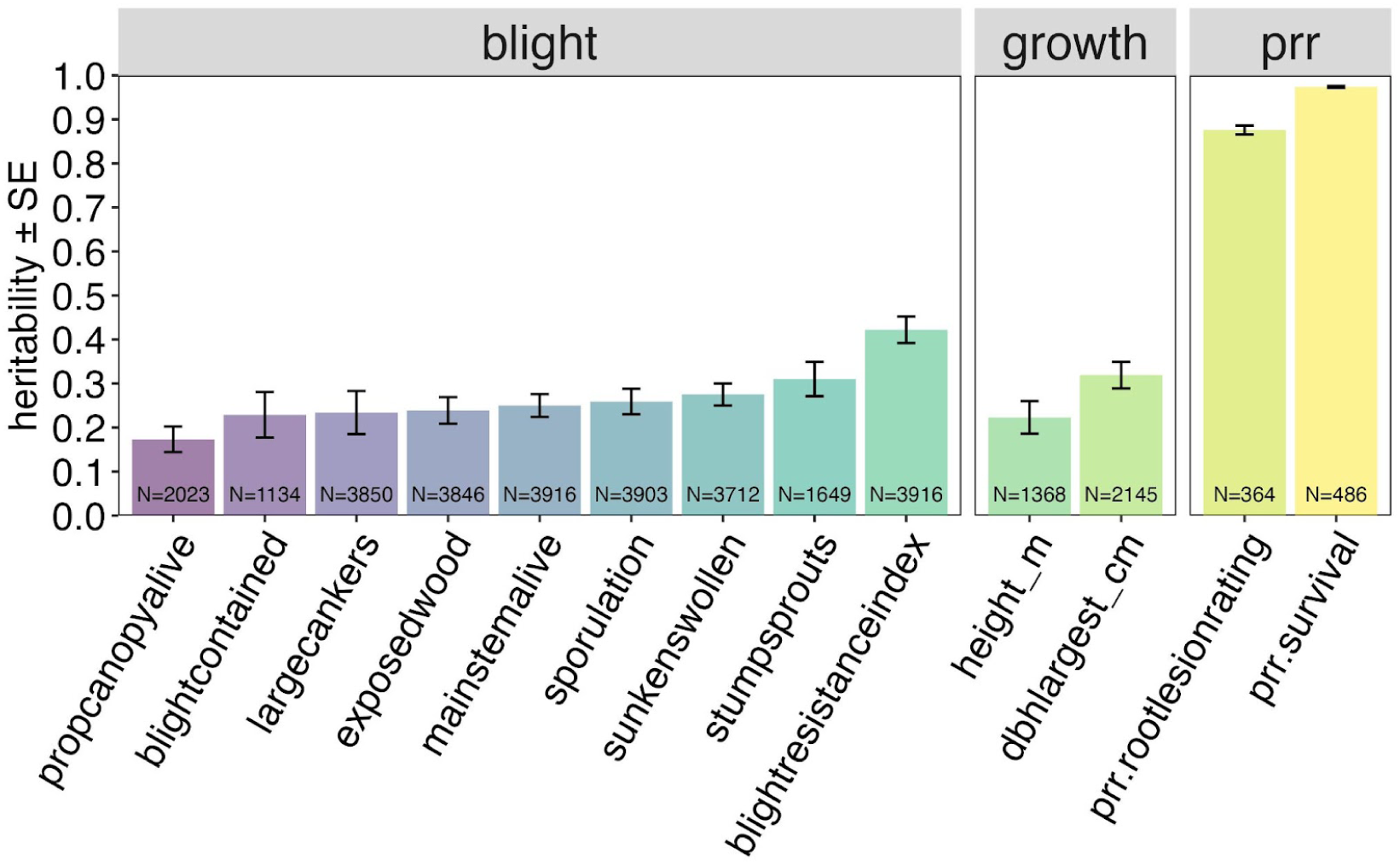
Heritability (h^2^) estimates for blight resistance, PRR resistance, and growth traits. Heritabilities were estimated from genomic relationships among the subset of genotyped and phenotyped trees. Numbers are sample sizes of trees used to estimate h^2^ for each trait.

**Fig. S2.**
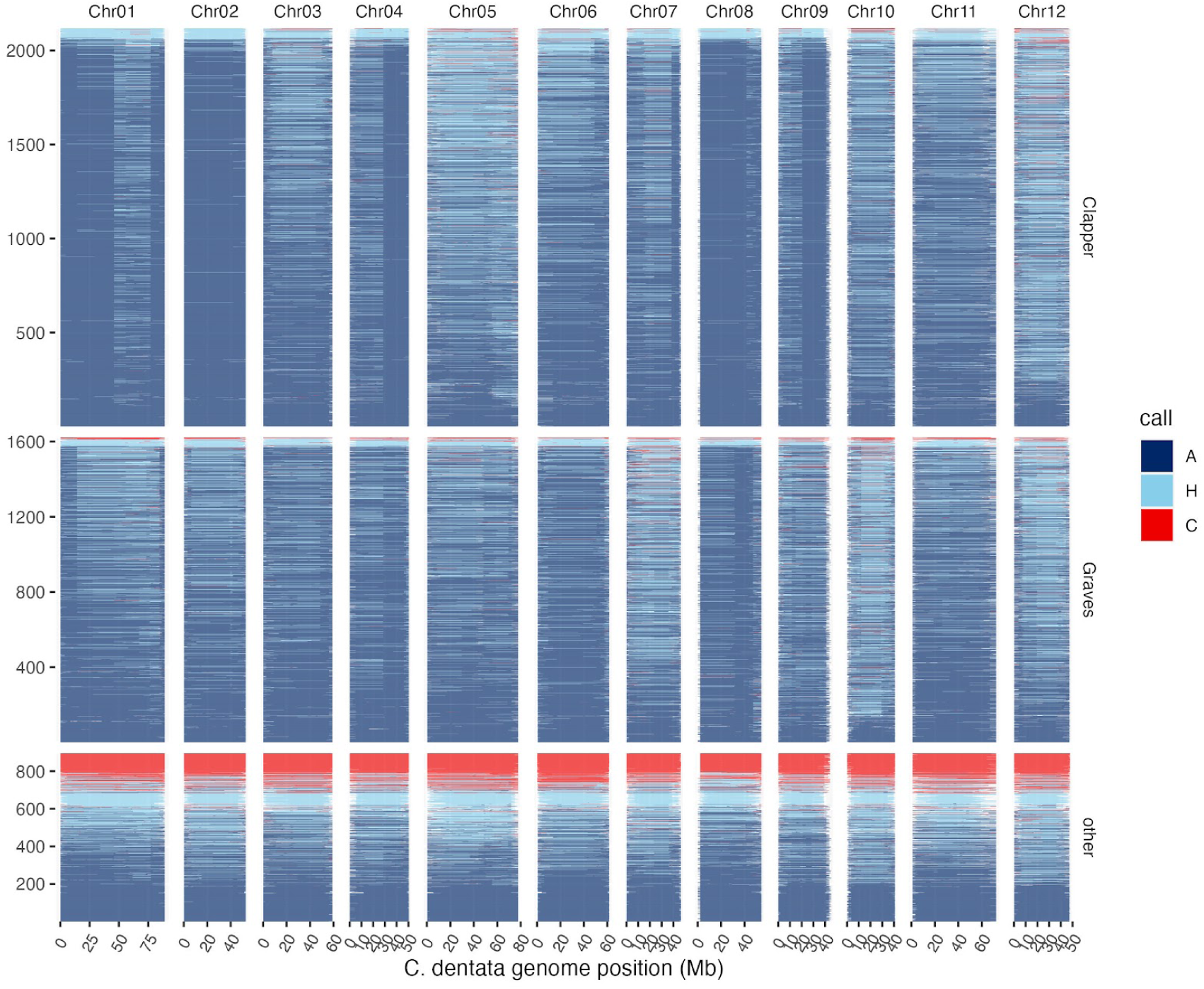
Local species ancestry among *C. dentata* backcross hybrids. Local ancestry calls were made for descendants of the Clapper BC_1_ tree, the Graves F_1_ tree, and other sources of resistance in Ancestry_HMM. Numbers of trees from each population are depicted on the y-axis. Genomic regions where trees inherited both alleles from *C. dentata* or *C. mollissima* are depicted as dark blue and red regions, respectively, and regions of heterozygous ancestry are depicted as light blue regions.

**Fig. S3.**
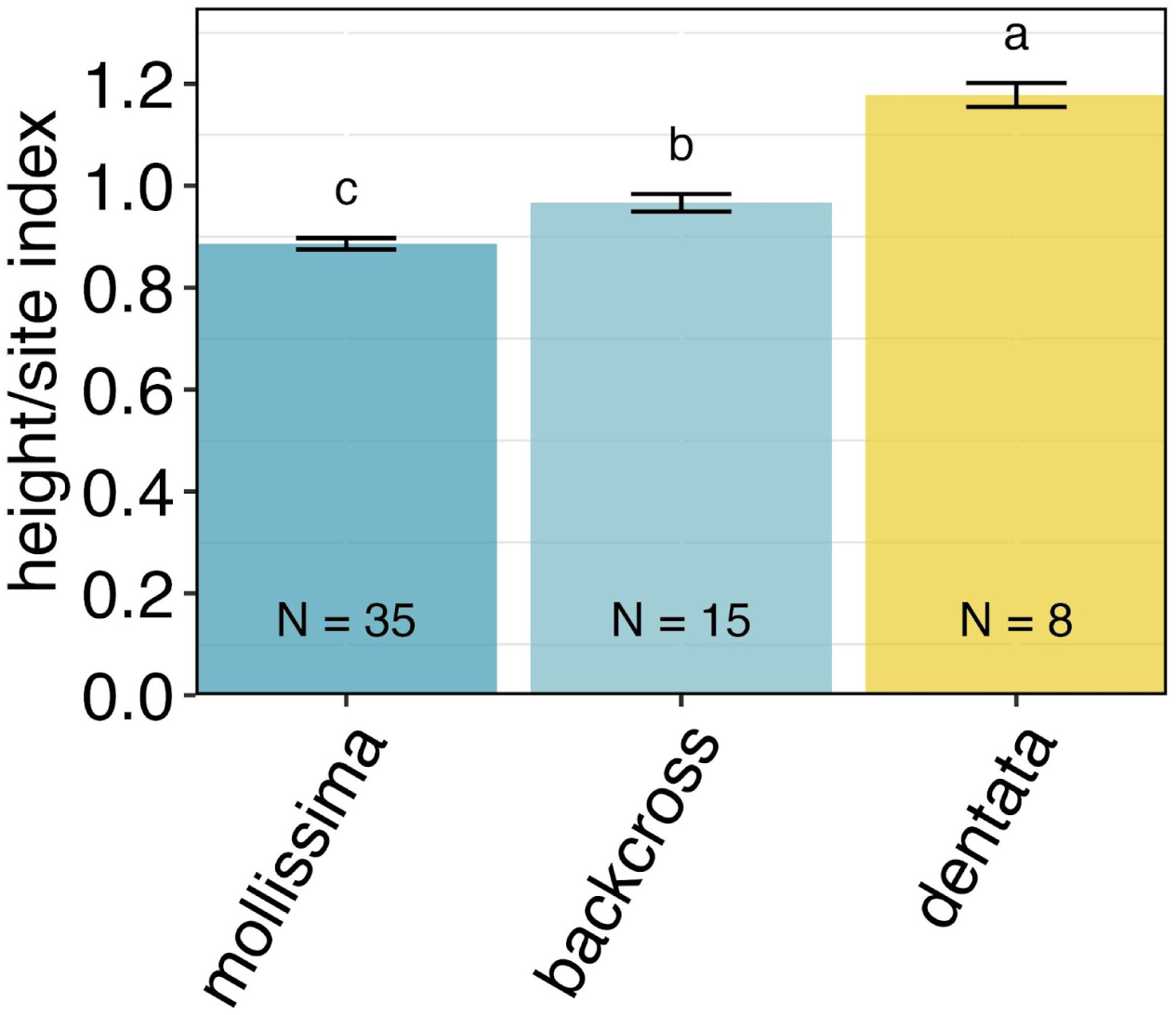
Comparison of tree heights for *C. mollissima*, *C. dentata* backcross, and large surviving American chestnut (LSAs) in a closed canopy stand at Lesesne State Forest, VA. Trees varied in age from 22 to 33 years old. To account for age-based differences in tree height, observed tree heights were divided by age-based expectations for yellow poplar (*Liriodendron tulipifera*) at site index = 90 (*124*). Age adjusted tree heights were compared for the 30% fastest growing individuals from each genotype class.

**Fig. S4.**
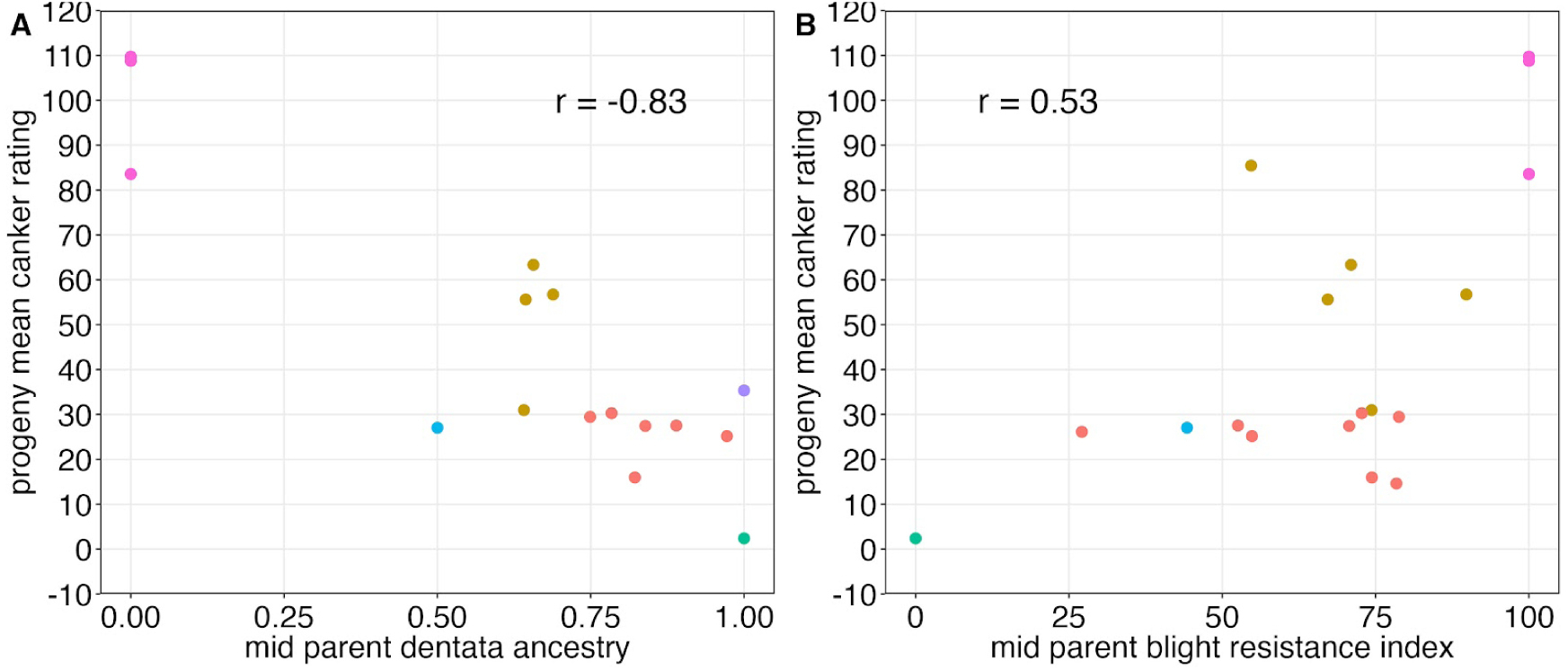
Correlations between progeny mean blight resistance and parental *C. dentata* ancestry (A) or blight resistance (B). Colors indicate different cross types (pink - *C. mollissima*, light blue - C. mollissima x C. dentata F_1_, green = *C. dentata,* purple - large surviving American chestnut, red - *C. dentata* BC_3_F_2_, brown - *C. dentata* backcross x F_1_). Only the subset of families where both parents were genotyped (A, 16 families) or phenotyped (B, 18 families) are shown. A summary of resistance and ancestry data for the entire population of 30 families can be found in Table S1.

**Fig. S5.**
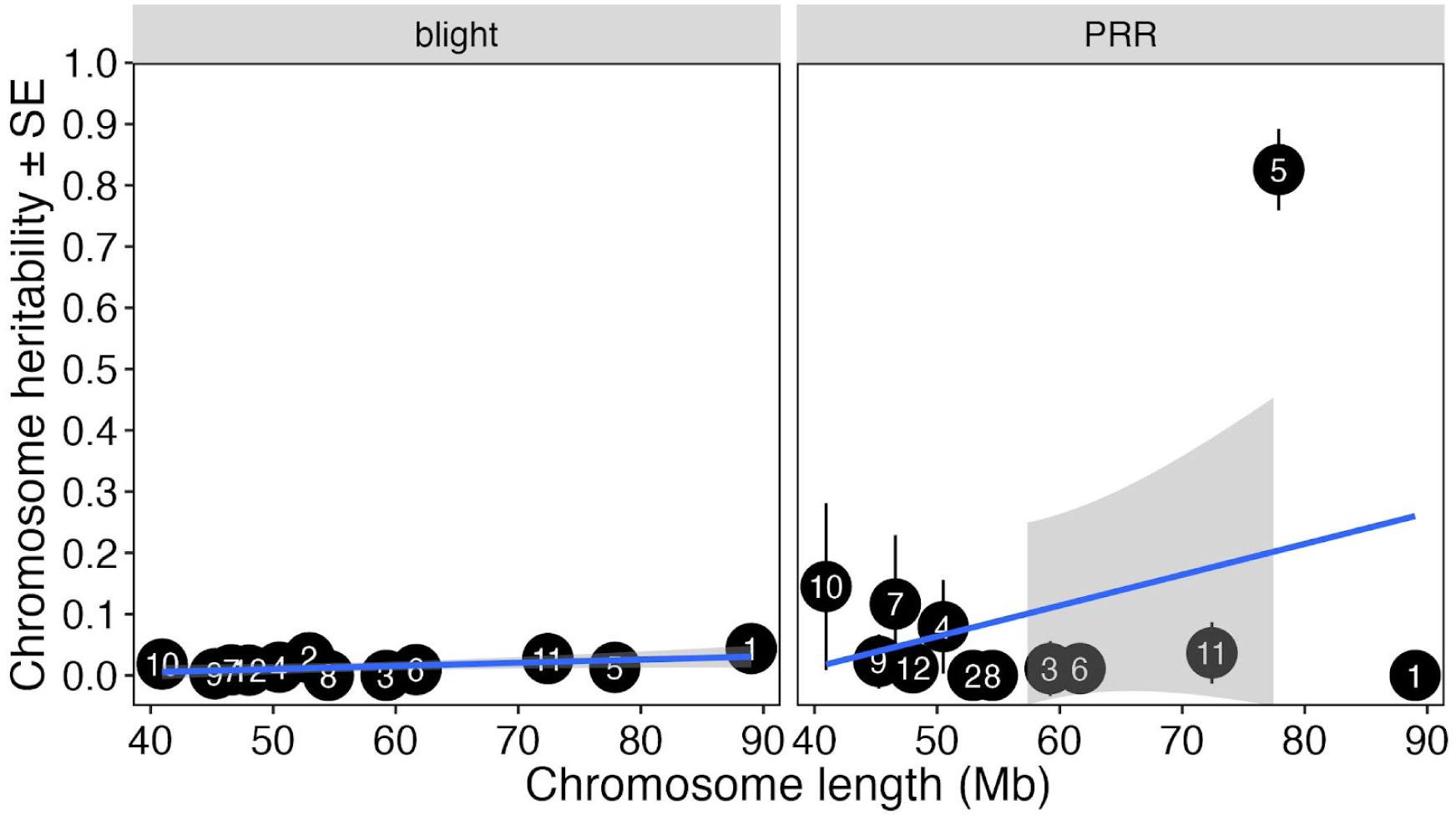
Proportion of phenotypic variance explained (chromosome heritability) versus chromosome length for the blight resistance index and PRR survival. Lines are the linear regression ± 95% confidence intervals. A summary of the chromosome heritability for blight and PRR resistance can be found in Table S2.

**Fig. S6.**
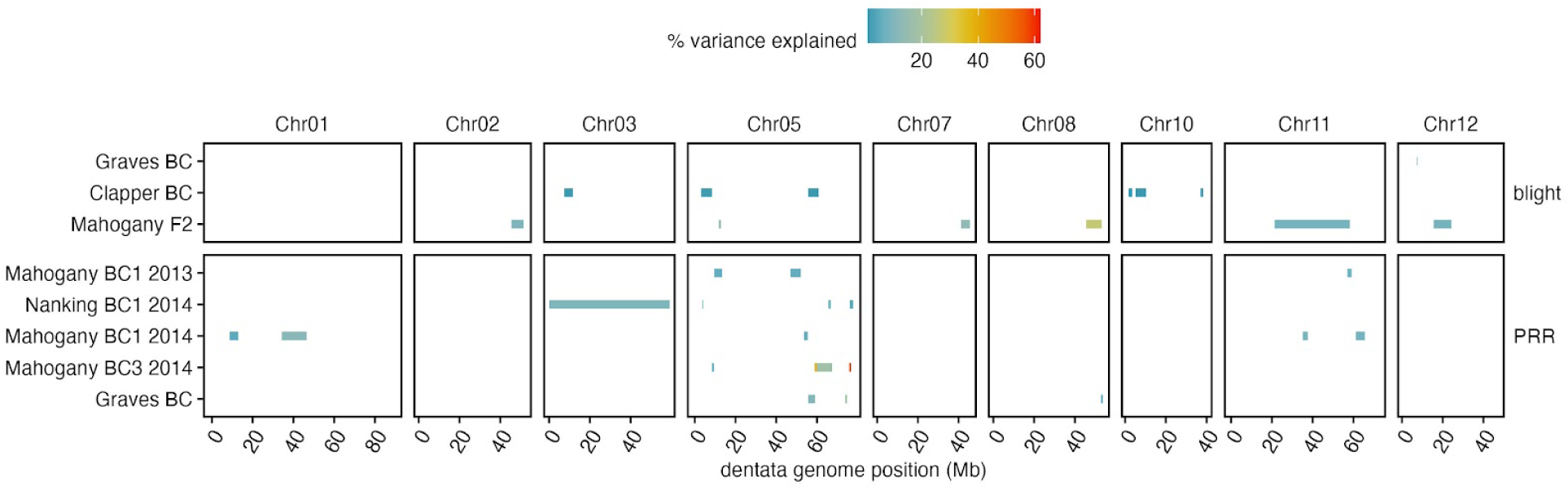
Comparisons of the locations of blight and PRR resistance QTL peaks across mapping populations. The Graves and Clapper backcross population QTLs mapped in the current study are compared to QTLs mapped in previous studies (*26*, *54*). Summary statistics for QTL locations and effect sizes can be found in the supplementary data.

**Fig. S7.**
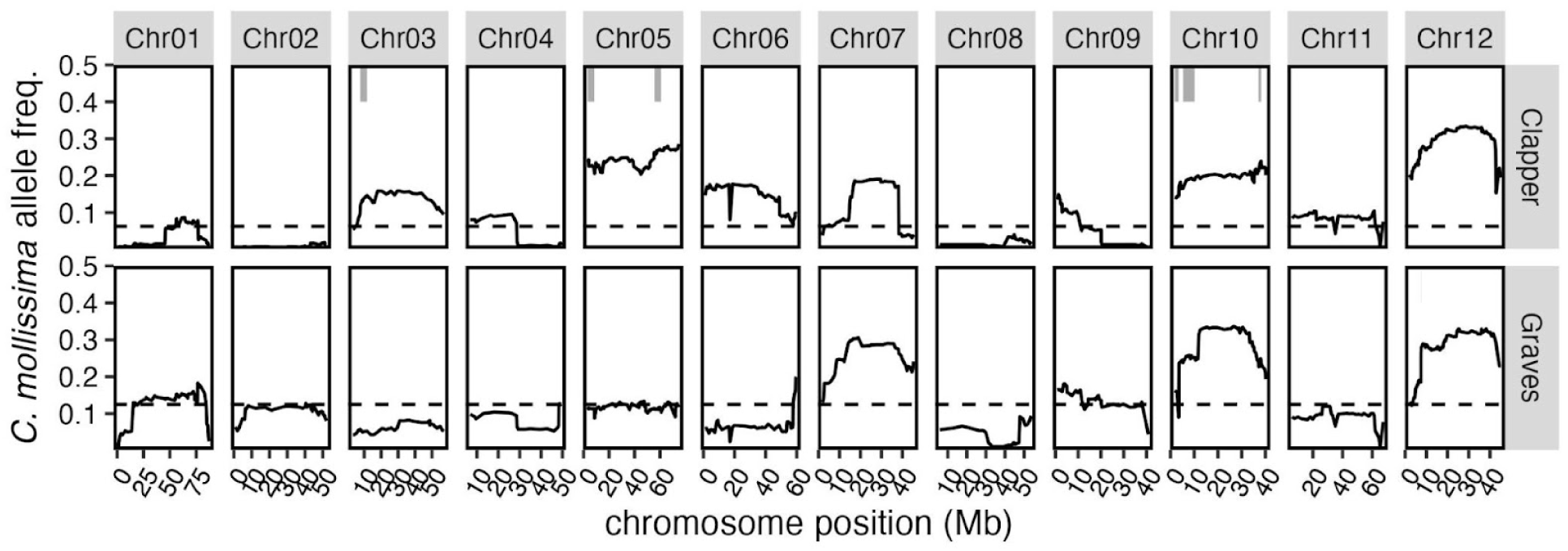
Frequency of *C. mollissima* alleles in among *C. dentata* backcross progeny. Local *C. mollissima* v. *C. dentata* ancestry for individual trees was inferred in Ancestry_HMM. Allele frequencies were calculated for 265 BC_3_F_2_ descendants of the ‘Clapper’ BC_1_ tree and 279 BC_2_F_2_ descendants of the ‘Graves’ F_1_ tree that had blight resistance index values > 50. Locations of blight resistance QTL intervals detected in each population are displayed as grey bars. Dotted lines depicted expected *C. mollissima* allele frequencies at BC_3_F_2_ (0.0625) and BC_2_F_2_ (0.125) in the absence of selection.

**Table S1.**
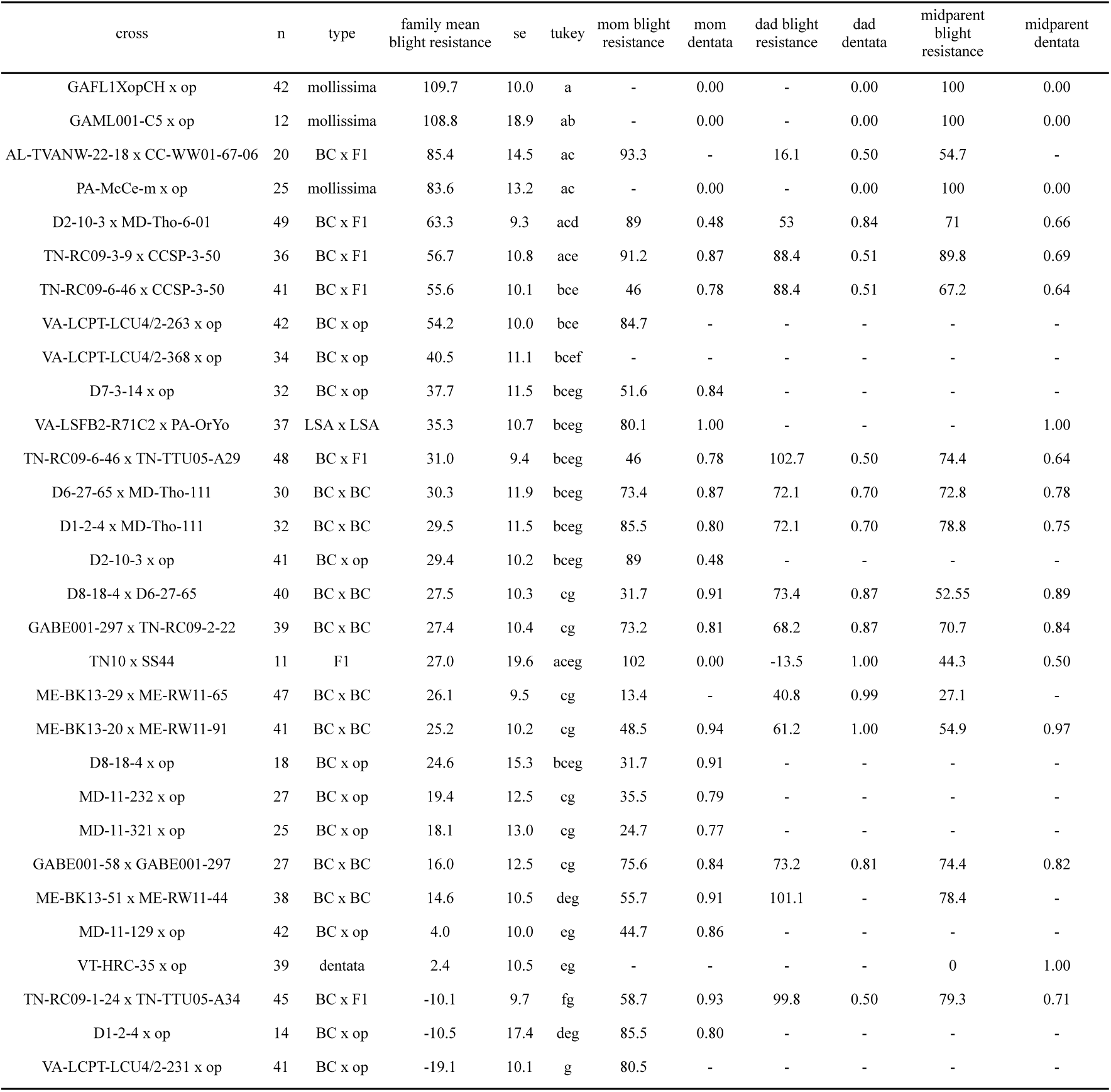
Comparison of progeny and parental blight resistance estimates mean blight resistance. Blight resistance ratings for seedling progeny were scaled from 0 = mean of susceptible *C. dentata* controls to 100 = mean of resistant *C. mollissima* controls and Tukey tests were performed for comparisons of family means. Progeny blight resistance was compared to estimates of parental blight resistance index and *C. dentata* ancestry proportions.

**Table S2.**
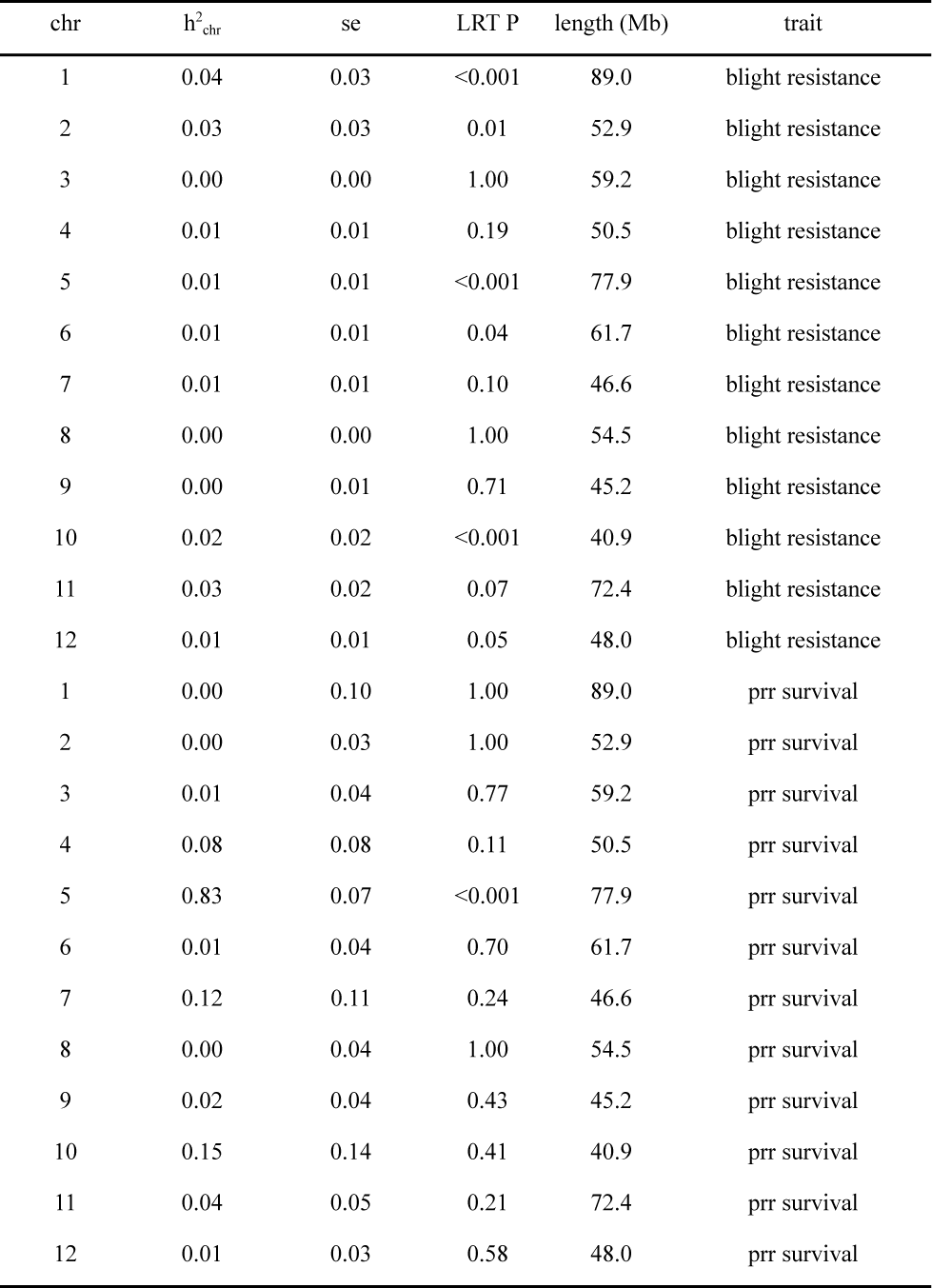
Heritability of blight and PRR resistance by chromosome. Proportion of phenotypic variance in blight resistance and PRR survival indices (h^2^_chr_) was estimated in ASReml by fitting models with two genomic relationship matrices 1. for the individual chromosomes and 2. the rest of the genome. Likelihood ratio tests (LRT) were performed to determine if individual chromosomes explain significant proportions of the variance in resistance. For both blight and PRR resistance, variance explained per chromosome was uncorrelated with chromosome length (Mb).

**Table S3.**
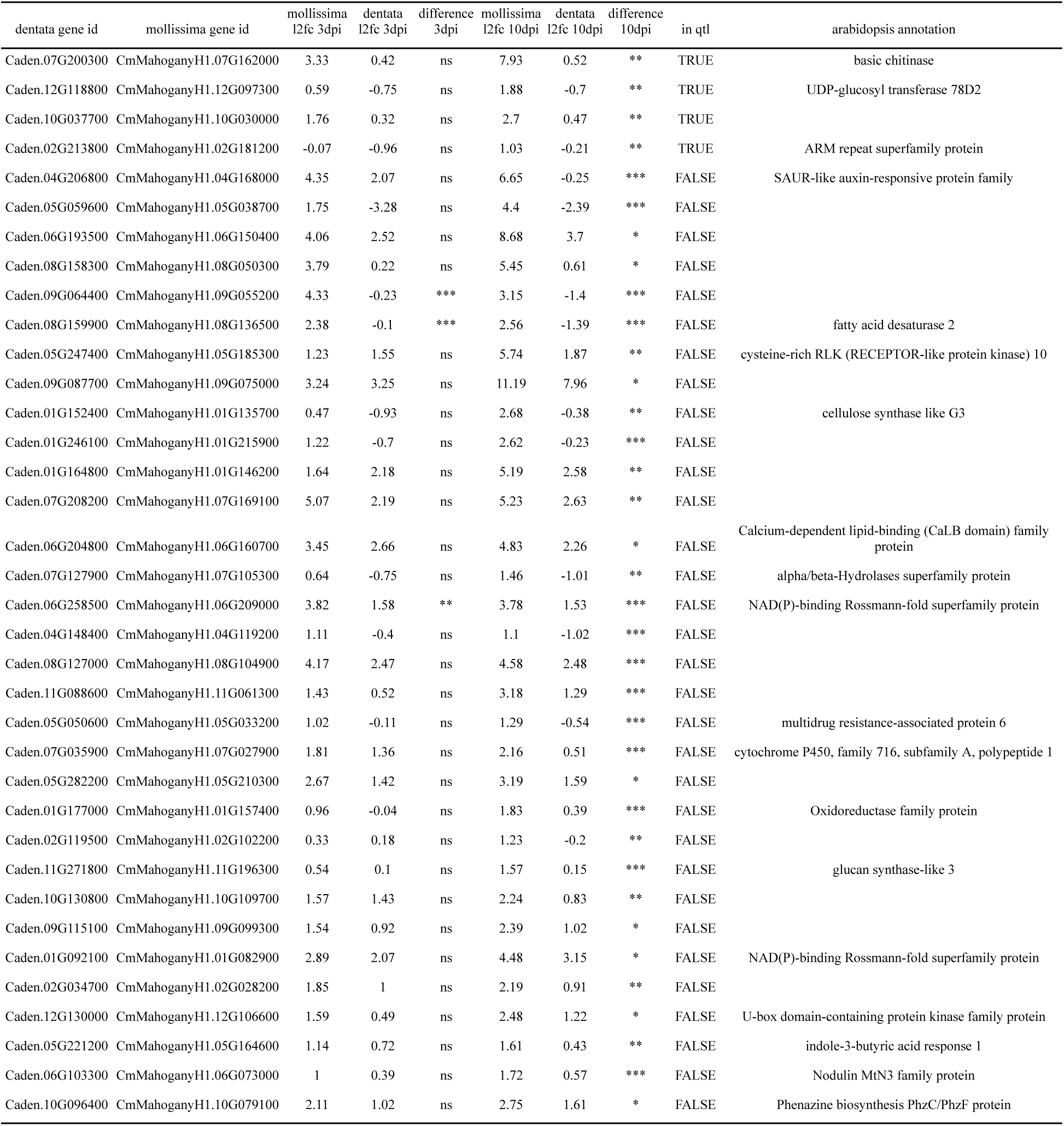
Genes with significant *C. mollissima* allele-specific upregulated expression in *C. dentata* x *C. mollissima* F_1_ hybrids in response to *C. parasitica* inoculation. Log 2 fold change (l2fc) in the expression of *C. dentata* and *C. mollissima* alleles in response to *C. parasitica* are reported for 3 day (3dpi) and 10 day (10dpi) post inoculation timepoints. Significant *C. mollissima* - *C. dentata* allele specific expression differences are reported for Benjamini-Hochberg false discovery rates ranging from 0.05 to 0.01 (*), from 0.01 to 0.001 (**), and less than 0.001 (***). Four genes mapped to blight QTLs (in qtl). The putative functions of these genes are reported from the annotation of *Arabidopsis thaliana* homologs.

**Table S4.**
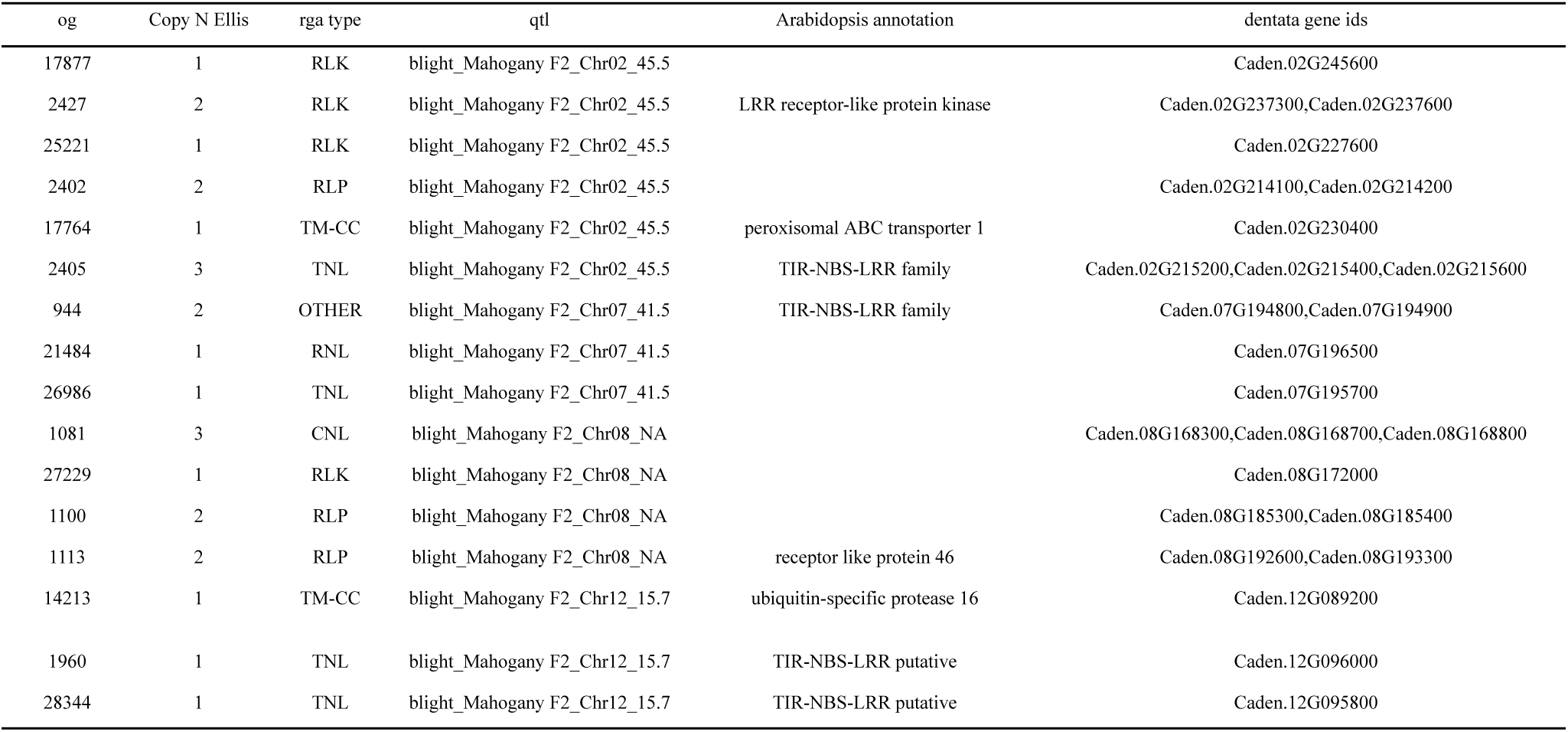
Resistance gene orthogroups that map to blight QTLs in a Mahogany F_2_ population *C. dentata* ‘Ellis’ and absent in both haplotypes of *C. mollissima* ‘Mahogany’. These genes are candidates for blight susceptibility in *C. dentata* under the hypothesis that necrotrophic fungi secrete effector proteins that interact with host ‘resistance’ proteins to trigger cell death. Copy numbers of genes with orthogroups (og) were compared using GENESPACE(*67*) and ‘resistance gene analog’ types were identified with RGAugury (*120*).

**Table S5.** *Cryphonectria parasitica* effectors with upregulated expression in *C. dentata*. Expression *in planta* was compared to expression *in vitro* in potato dextrose media. Log2 fold change (l2fc) and Benjamini-Hochberg false discovery rates (padj) are reported at 3 days and 10 days post inoculation (dpi). EffectorP v3 (*121*) was used to classify effectors as cytoplasmic and/or apoplastic. Gene ids and annotation are from v2 of the C. *parasitica* genome (*122*).

| gene | effectorp | 3d basemean | 3d l2fc | 3d padj | 10d basemean | 10d l2fc | 10 d padj | annotation |
| --- | --- | --- | --- | --- | --- | --- | --- | --- |
| 354471 | Apoplastic | 255.92 | 8.28 | 0.01 | 1728.44 | 9.91 | 0 | - |
| 327572 | Apoplastic | 95.64 | 8.93 | 0.01 | 181.69 | 9.05 | 0 | - |
| 47319 | Apoplastic | 1.82 | 5.82 | - | 40.44 | 8.95 | 0 | - |
| 67264 | Apoplastic | 70.4 | 8.47 | 0 | 110.6 | 8.37 | 0 | Polygalacturonase |
| 223321 | Apoplastic | 0.59 | 5.72 | - | 7.53 | 7.52 | 0.03 | - |
| 75076 | Apoplastic | 0.67 | 4.63 | - | 21.19 | 7.46 | 0 | - |
| 39058 | Apoplastic | 39.01 | 4.96 | 0.06 | 374.35 | 7.26 | 0 | - |
| 255494 | Apoplastic | 3.09 | 5.1 | 0.15 | 22.63 | 6.86 | 0 | - |
| 337842 | Apoplastic | 1.78 | 5.12 | - | 8.24 | 6.43 | 0.03 | - |
| 326343 | Apoplastic | 13.85 | 5.15 | 0 | 53.44 | 6.19 | 0 | - |
| 331841 | Apoplastic | 17.33 | 7.14 | 0 | 11.19 | 6 | 0.04 | - |
| 51476 | Apoplastic | 21.11 | 7.14 | 0 | 8.49 | 5.51 | 0.14 | Polygalacturonase |
| 351651 | Apoplastic | 9.57 | 4.73 | 0.04 | 31.18 | 5.44 | 0 | - |
| 38857 | Apoplastic | 3.64 | 3.07 | 0.67 | 35.55 | 5.38 | 0 | - |
| 49953 | Apoplastic | 128.95 | 6.37 | 0.08 | 100.8 | 5.35 | 0 | - |
| 60731 | Apoplastic | 4.39 | 3.84 | 0.22 | 17.37 | 5 | 0.03 | Galacturan 1,4-alpha-galacturonidase |
| 331312 | Apoplastic | 76.83 | 6.5 | 0.13 | 34.08 | 4.77 | 0.01 | - |
| 29438 | Apoplastic | 81.82 | 5.35 | 0.13 | 65.07 | 4.34 | 0.02 | - |
| 19334 | Apoplastic | 30.13 | 4.6 | 0.19 | 39.57 | 4.26 | 0.02 | - |
| 62910 | Apoplastic | 37.96 | 6.05 | 0.05 | 37.48 | 5.17 | 0.12 | - |
| 71339 | Apoplastic/cytoplasmic | 143.82 | 5.20 | 0.26 | 114.87 | 4.09 | 0.05 | - |
| 74551 | Apoplastic | 9.11 | 2.32 | 0.99 | 57.64 | 4.21 | 0.02 | - |

## DESCRIPTION OF SUPPLEMENTARY DATA

Accessible on Dryad: https://doi.org/10.5061/dryad.4xgxd25mj

**Supplementary Data 1 | RNA-seq metadata**. Contains species, tissue, and treatment information for samples used for genome annotation and *C. parasitica* gene expression responses.

**Supplementary Data 2 | Combined bed-like annotation file**. This contains syntenic orthogroup information, as produced by GENESPACE. The standard bed file with a set of additional fields presenting orthogroup and tandem array information for each gene.

**Supplementary Data 3 | GENESPACE pan-gene sets against the Ellis coordinate system.** Pangenes file presenting the interpolated syntenic position of each syntenic phylogenetically hierarchical orthogroup.

**Supplementary Data 4 | Syntenic block breakpoints.** Syntenic blocks were inferred with DEEPSPACE using default parameters except as noted in the methods. This is a standard .paf file except that the first field contains the DEEPSPACE file identifier and the last two fields contain the identifier for the query (windows) and target genomes.

**Supplementary Data 5 | Blight and growth phenotypes.** Blight presence/absence phenotypes were coded into 0 (susceptible) and 1 (resistant) phenotypic classes. File includes tree inoculation, status, tree age, climate, and soil variables relevant for *C. dentata* habitat suitability that were modeled as fixed effects.

**Supplementary Data 6 | PRR resistance phenotypes.** Survival and root lesion severity data after *Phytophthora cinnamomi* inoculation for ∼27 k open pollinated progeny of *C. dentata* backcross hybrids, resistant *C. mollissima,* & susceptible *C. dentata* controls.

**Supplementary Data 7 | Multi-generation pedigree for all trees phenotyped in this study.** Includes maternal and paternal parents for multiple generations of hybrid trees in The American Chestnut Foundation breeding program.

**Supplementary Data 8 | Genotyping-by-sequencing data from 5,003 trees in variant calling format (VCF).**

**Supplementary Data 9 | Genetic maps for the ‘GMBig x Horn’ *C. dentata* full sib family.** Contains genetic map positions in centiMorgans and physical coordinates in the ‘Ellis’ genome for each parent from this cross.

**Supplementary Data 10 | Hybrid ancestry estimates.** Hybrid species ancestry inferences for ∼5 k trees from Ancestry_HMM.

**Supplementary Data 11 | Estimated breeding values for blight resistance, PRR resistance, and tree size.** Breeding values for blight and PRR resistance indices that were scaled from 0 (*C. dentata* mean) to 100 (*C. mollissima* mean). Resistance data were merged with hybrid species ancestry estimates from Ancestry_HMM

**Supplementary Data 12 | Progeny blight resistance phenotypic data.** Seedling blight resistance phenotypes for ∼1 K progeny from 30 full or half sib families

**Supplementary Data 13 | Blight and PRR resistance QTL interval metadata.** Peak and border coordinates and effect sizes for blight and PRR QTL intervals across the *C. dentata* and *C. mollissima* genomes.

**Supplementary Data 14 | Species and allele specific expression responses to *C. parasitica*.** Log_2_fold change and P-values for species- and allele-specific expression responses to *C. parasitica* inoculation.

**Supplementary Data 15 | Resistance gene analogs (RGAs) in the *C. dentata* and *C. mollissima* genomes.** Contains output from RGAugury to classify resistance genes across *C. dentata* ‘Ellis’ and *C. mollissima* ‘Mahogany’ and ‘Nanking’ reference genomes.

**Supplementary Data 16 | Comparison of *C. parasitica* gene expression responses in *C. mollissima* and *C. dentata* hosts.** Contains log2 fold change and P-values for the expression of C. parasitica genes in resistant C. mollissima versus susceptible C. dentata hosts. Effectors from *C. parasitica* were identified using EffectorP (v3).

## Notes

### Competing Interest Statement

The authors have declared no competing interest.

## REFERENCES

1. S. L. Anagnostakis, Chestnut Blight: The Classical Problem of an Introduced Pathogen. Mycologia 79, 23–37 (1987).

2. C. L. Shear, N. E. Stevens, The chestnut-blight parasite (*Endothia parasitica)* from China. Science 38, 295–297 (1913).

3. F. V. Hebard, Developmental histopathology of cankers incited by hypovirulent and virulent isolates of*Endothia parasitica*on susceptible and resistant chestnut trees. Phytopathology 74, 140 (1984).

4. D. E. Davis, The American Chestnut: An Environmental History (University of Georgia Press, 2021).

5. H. J. Dalgleish, C. D. Nelson, J. A. Scrivani, D. F. Jacobs, Consequences of shifts in abundance and distribution of American chestnut for restoration of a foundation forest tree. For. Trees Livelihoods 7, 4 (2015).

6. K. M. Potter, B. S. Crane, W. W. Hargrove, A United States national prioritization framework for tree species vulnerability to climate change. New Forests 48, 275–300 (2017).

7. A. M. Sandercock, J. W. Westbrook, Q. Zhang, J. A. Holliday, A genome-guided strategy for climate resilience in American chestnut restoration populations. Proc. Natl. Acad. Sci. U. S. A. 121, e2403505121 (2024).

8. S. L. Anagnostakis, Chestnut breeding in the United States for disease and insect resistance. Plant Dis. 96, 1392–1403 (2012).

9. B. S. Crandall, G. F. Gravatt, M. Ryan, Root disease of *Castanea* species and some coniferous and broadleaf nursery stocks, caused by *Phytophthora cinnamomi*. Phytopathology 35 (1945).

10. T. I. Burgess, J. K. Scott, K. L. Mcdougall, M. J. C. Stukely, C. Crane, W. A. Dunstan, F. Brigg, V. Andjic, D. White, T. Rudman, F. Arentz, N. Ota, G. E. S. J. Hardy, Current and projected global distribution of *Phytophthora cinnamomi*, one of the world’s worst plant pathogens. Glob. Chang. Biol. 23, 1661–1674 (2017).

11. K. C. Steiner, J. W. Westbrook, F. V. Hebard, L. L. Georgi, W. A. Powell, S. F. Fitzsimmons, Rescue of American chestnut with extraspecific genes following its destruction by a naturalized pathogen. New Forests 48, 317–336 (2017).

12. H. J. Dalgleish, R. K. Swihart, American chestnut past and future: implications of restoration for resource pulses and consumer populations of eastern US forests. Restor. Ecol., doi: 10.1111/j.1526-100X.2011.00795.x.

13. J. G. Skousen, K. Dallaire, S. Scagline-Mellor, A. Monteleone, L. Wilson-Kokes, J. Joyce, C. Thomas, T. Keene, C. DeLong, T. Cook, D. F. Jacobs, Plantation performance of chestnut hybrids and progenitors on reclaimed Appalachian surface mines. New For (Dordr) 49, 599–611 (2018).

14. E. J. Gustafson, B. R. Sturtevant, A. M. G. de Bruijn, N. Lichti, D. F. Jacobs, D. M. Kashian, B. R. Miranda, P. A. Townsend, Forecasting effects of tree species reintroduction strategies on carbon stocks in a future without historical analog. Glob. Chang. Biol. 24, 5500–5517 (2018).

15. S. L. Clark, S. E. Schlarbaum, A. M. Saxton, S. N. Jeffers, R. E. Baird, Eight-year field performance of backcross American chestnut (*Castanea dentata*) seedlings planted in the southern Appalachians, USA. For. Ecol. Manage. 532, 120820 (2023).

16. G. J. Griffin, F. V. Hebard, R. W. Wendt, J. R. Elkins, Others, Survival of American chestnut trees: evaluation of blight resistance and virulence in *Endothia parasitica*. Phytopathology 73, 1084–1092 (1983).

17. G. J. Griffin, J. R. Elkins, D. McCurdy, S. L. Griffin, Integrated use of resistance, hypovirulence, and forest management to control blight on American chestnut. Restoration of American chestnut to forest lands, 97–107 (2006).

18. A. H. Graves, Relative Blight Resistance in Species and Hybrids of Castanea (1950).

19. G. F. Gravatt, J. D. Diller, F. H. Berry, A. H. Graves, H. Nienstaedt, Breeding timber chestnuts for blight resistance in Northeastern Forest Tree Improvement Conference Proceedings (1953) vol. 1, pp. 70–75.

20. J. D. Diller, R. B. Clapper, Asiatic and hybrid chestnuts in the Eastern United States. Journal of Forestry 67, 328–331 (1969).

21. T. L. Kubisiak, F. V. Hebard, C. D. Nelson, J. Zhang, R. Bernatzky, H. Huang, S. L. Anagnostakis, R. L. Doudrick, Molecular mapping of resistance to blight in an interspecific cross in the genus *Castanea*. Phytopathology 87, 751–759 (1997).

22. C. R. Burnham, P. A. Rutter, D. W. French, Others, Breeding blight-resistant chestnuts. Plant Breed. Rev. 4, 347–397 (1986).

23. M. Diskin, K. C. Steiner, F. V. Hebard, Recovery of American chestnut characteristics following hybridization and backcross breeding to restore blight-ravaged Castanea dentata. For. Ecol. Manage. 223, 439–447 (2006).

24. J. W. Westbrook, J. B. James, P. H. Sisco, J. Frampton, S. Lucas, S. N. Jeffers, Resistance to *Phytophthora cinnamomi* in American chestnut (*Castanea dentata*) backcross populations that descended from two Chinese chestnut (*Castanea mollissima*) sources of resistance. Plant Dis. 103, 1631–1641 (2019).

25. J. W. Westbrook, Q. Zhang, M. K. Mandal, E. V. Jenkins, L. E. Barth, J. W. Jenkins, J. Grimwood, J. Schmutz, J. A. Holliday, Optimizing genomic selection for blight resistance in American chestnut backcross populations: A trade-off with American chestnut ancestry implies resistance is polygenic. Evol. Appl. 13, 31–47 (2020).

26. S. Fan, L. Georgi, F. Hebard, T. Zhebentyayeva, J. Yu, P. Sisco, S. Fitzsimmons, M. Staton, A. Abbott, C. Nelson, Mapping QTLs for blight resistance and morphological traits in inter-species hybrid families of chestnut (*Castanea spp*.). Front. Plant Sci. 15 (2024).

27. W. A. Powell, A. E. Newhouse, V. Coffey, Developing blight-tolerant American chestnut trees. Cold Spring Harb. Perspect. Biol. 11 (2019).

28. W. Powell, Petition for determination of nonregulated status for blight-tolerant darling 58 American chestnut. https://www.esf.edu/chestnut/documents/petition_executive_summary.pdf.

29. L. Bianco, P. Fontana, A. Marchesini, S. Torre, M. Moser, S. Piazza, S. Alessandri, V. Pavese, P. Pollegioni, C. Vernesi, M. Malnoy, D. Torello Marinoni, S. Murolo, L. Dondini, C. Mattioni, R. Botta, F. Sebastiani, D. Micheletti, L. Palmieri, The de novo, chromosome-level genome assembly of the sweet chestnut (*Castanea sativa* Mill.) Cv. Marrone Di Chiusa Pesio. BMC Genom Data 25, 64 (2024).

30. K. Shirasawa, S. Nishio, S. Terakami, R. Botta, D. T. Marinoni, S. Isobe, Chromosome-level genome assembly of Japanese chestnut (*Castanea crenata* Sieb. et Zucc.) reveals conserved chromosomal segments in woody rosids. DNA Res. 28 (2021).

31. Y. Xing, Y. Liu, Q. Zhang, X. Nie, Y. Sun, Z. Zhang, H. Li, K. Fang, G. Wang, H. Huang, T. Bisseling, Q. Cao, L. Qin, Hybrid de novo genome assembly of Chinese chestnut (*Castanea mollissima*). Gigascience 8 (2019).

32. J. Wang, P. Hong, Q. Qiao, D. Zhu, L. Zhang, K. Lin, S. Sun, S. Jiang, B. Shen, S. Zhang, Q. Liu, Chromosome-level genome assembly provides new insights into Japanese chestnut (*Castanea crenata*) genomes. Front. Plant Sci. 13, 1049253 (2022).

33. Y. Sun, Z. Lu, X. Zhu, H. Ma, Genomic basis of homoploid hybrid speciation within chestnut trees. Nat. Commun. 11, 3375 (2020).

34. M. Staton, C. Addo-Quaye, N. Cannon, J. Yu, T. Zhebentyayeva, M. Huff, N. Islam-Faridi, S. Fan, L. L. Georgi, C. D. Nelson, E. Bellis, S. Fitzsimmons, N. Henry, D. Drautz-Moses, R. E. Noorai, S. Ficklin, C. Saski, M. Mandal, T. K. Wagner, N. Zembower, C. Bodénès, J. Holliday, J. Westbrook, J. Lasky, F. V. Hebard, S. C. Schuster, A. G. Abbott, J. E. Carlson, A reference genome assembly and adaptive trait analysis of *Castanea mollissima* ‘Vanuxem,’ a source of resistance to chestnut blight in restoration breeding. Tree Genet. Genomes 16, 57 (2020).

35. B.-F. Zhou, S. Yuan, A. A. Crowl, Y.-Y. Liang, Y. Shi, X.-Y. Chen, Q.-Q. An, M. Kang, P. S. Manos, B. Wang, Phylogenomic analyses highlight innovation and introgression in the continental radiations of Fagaceae across the Northern Hemisphere. Nat. Commun. 13, 1320 (2022).

36. D. A. Larson, M. E. Staton, B. Kapoor, N. Islam-Faridi, T. Zhebentyayeva, S. Fan, J. Stork, A. Thomas, A. S. Ahmed, E. C. Stanton, A. Houston, S. E. Schlarbaum, M. W. Hahn, J. E. Carlson, A. G. Abbott, S. DeBolt, C. Dana Nelson, A haplotype-resolved reference genome of Quercus alba sheds light on the evolutionary history of oaks. New Phytologist (2024).

37. G. Bazzigher, G. A. Miller, Blight-resistant chestnut selections of Switzerland : A valuable germplasm resource. Plant Disease 75, 5–9 (1987).

38. T. H. Meuwissen, B. J. Hayes, M. E. Goddard, Prediction of total genetic value using genome-wide dense marker maps. Genetics 157, 1819–1829 (2001).

39. D. Grattapaglia, O. B. Silva-Junior, R. T. Resende, E. P. Cappa, B. S. F. Müller, B. Tan, F. Isik, B. Ratcliffe, Y. A. El-Kassaby, Quantitative genetics and genomics converge to accelerate forest tree breeding. Front. Plant Sci. 9, 1693 (2018).

40. D. Habier, R. L. Fernando, J. C. M. Dekkers, Genomic selection using low-density marker panels. Genetics 182, 343–353 (2009).

41. D. Hall, H. R. Hallingbäck, H. X. Wu, Estimation of number and size of QTL effects in forest tree traits. Tree Genet. Genomes 12 (2016).

42. J. Yang, T. A. Manolio, L. R. Pasquale, E. Boerwinkle, N. Caporaso, J. M. Cunningham, M. de Andrade, B. Feenstra, E. Feingold, M. G. Hayes, W. G. Hill, M. T. Landi, A. Alonso, G. Lettre, P. Lin, H. Ling, W. Lowe, R. A. Mathias, M. Melbye, E. Pugh, M. C. Cornelis, B. S. Weir, M. E. Goddard, P. M. Visscher, Genome partitioning of genetic variation for complex traits using common SNPs. Nat. Genet. 43, 519–525 (2011).

43. F. L. Paillet, P. A. Rutter, Replacement of native oak and hickory tree species by the introduced American chestnut (*Castanea dentata*) in southwestern Wisconsin. Can. J. Bot. 67, 3457–3469 (1989).

44. K. Baier, C. Maynard, W. Powell, Chestnuts and light: Early flowering in chestnut species induced under high-intensity, high-dose light in growth chambers. Journal of the American Chestnut Foundation 26, 8–10 (2012).

45. T. Mackay, E. A. Stone, J. Ayroles, The genetics of quantitative traits: challenges and prospects. Nat. Rev. Genet. 10, 565–577 (2009).

46. P. J. Wittkopp, G. Kalay, Cis-regulatory elements: molecular mechanisms and evolutionary processes underlying divergence. Nat. Rev. Genet. 13, 59–69 (2011).

47. M. Kumar, A. Brar, M. Yadav, A. Chawade, V. Vivekanand, N. Pareek, Chitinases—potential candidates for enhanced plant resistance towards fungal pathogens. Agriculture 8, 88 (2018).

48. D. Shao, D. L. Smith, M. Kabbage, M. G. Roth, Effectors of plant necrotrophic fungi. Front. Plant Sci. 12, 687713 (2021).

49. O. X. Dong, P. C. Ronald, Genetic engineering for disease resistance in plants: Recent progress and future perspectives. Plant Physiol. 180, 26–38 (2019).

50. H. Garcia-Ruiz, B. Szurek, G. Van den Ackerveken, Stop helping pathogens: engineering plant susceptibility genes for durable resistance. Curr. Opin. Biotechnol. 70, 187–195 (2021).

51. Z. Li, S. Parris, C. A. Saski, A simple plant high-molecular-weight DNA extraction method suitable for single-molecule technologies. Plant Methods 16, 38 (2020).

52. C.-L. Xiao, Y. Chen, S.-Q. Xie, K.-N. Chen, Y. Wang, Y. Han, F. Luo, Z. Xie, MECAT: fast mapping, error correction, and de novo assembly for single-molecule sequencing reads. Nat. Methods 14, 1072–1074 (2017).

53. C.-S. Chin, D. H. Alexander, P. Marks, A. A. Klammer, J. Drake, C. Heiner, A. Clum, A. Copeland, J. Huddleston, E. E. Eichler, S. W. Turner, J. Korlach, Nonhybrid, finished microbial genome assemblies from long-read SMRT sequencing data. Nat. Methods 10, 563–569 (2013).

54. T. N. Zhebentyayeva, P. H. Sisco, L. L. Georgi, S. N. Jeffers, M. T. Perkins, J. B. James, F. V. Hebard, C. Saski, C. D. Nelson, A. G. Abbott, Dissecting resistance to *Phytophthora cinnamomi* in interspecific hybrid chestnut crosses using sequence-based genotyping and QTL mapping. Phytopathology 109, 1594–1604 (2019).

55. N. C. Durand, M. S. Shamim, I. Machol, S. S. P. Rao, M. H. Huntley, E. S. Lander, E. L. Aiden, Juicer provides a one-click system for analyzing loop-resolution hi-C experiments. Cell Syst. 3, 95–98 (2016).

56. H. Cheng, G. T. Concepcion, X. Feng, H. Zhang, H. Li, Haplotype-resolved de novo assembly using phased assembly graphs with hifiasm. Nat. Methods 18, 170–175 (2021).

57. R. Vaser, I. Sović, N. Nagarajan, M. Šikić, Fast and accurate de novo genome assembly from long uncorrected reads. Genome Res. 27, 737–746 (2017).

58. S. Shu, D. Goodstein, D. Rokhsar, PERTRAN: Genome-Guided RNA-Seq Read Assembler (2013; https://www.osti.gov/biblio/1241180).

59. T. D. Wu, S. Nacu, Fast and SNP-tolerant detection of complex variants and splicing in short reads. Bioinformatics 26, 873–881 (2010).

60. J. T. Lovell, J. Jenkins, D. B. Lowry, S. Mamidi, A. Sreedasyam, X. Weng, K. Barry, J. Bonnette, B. Campitelli, C. Daum, S. P. Gordon, B. A. Gould, A. Khasanova, A. Lipzen, A. MacQueen, J. D. Palacio-Mejía, C. Plott, E. V. Shakirov, S. Shu, Y. Yoshinaga, M. Zane, D. Kudrna, J. D. Talag, D. Rokhsar, J. Grimwood, J. Schmutz, T. E. Juenger, The genomic landscape of molecular responses to natural drought stress in Panicum hallii. Nat. Commun. 9, 5213 (2018).

61. B. J. Haas, A. L. Delcher, S. M. Mount, J. R. Wortman, R. K. Smith, L. I. Hannick, R. Maiti, C. M. Ronning, D. B. Rusch, C. D. Town, S. L. Salzberg, O. White, Improving the Arabidopsis genome annotation using maximal transcript alignment assemblies. Nucleic Acids Res., doi: 10.1093/nar/gkg770 (2003).

62. A. F. A. Smit, Smit, AFA, Hubley, R & Green, P. RepeatMasker Open-3.0. [Preprint] (2015).

63. W. Bao, K. K. Kojima, O. Kohany, Repbase Update, a database of repetitive elements in eukaryotic genomes. Mob. DNA 6, 11 (2015).

64. A. A. Salamov, V. V. Solovyev, Ab initio gene finding in Drosophila genomic DNA. Genome Res., doi: 10.1101/gr.10.4.516 (2000).

65. G. S. C. Slater, E. Birney, Automated generation of heuristics for biological sequence comparison. BMC Bioinformatics, doi: 10.1186/1471-2105-6-31 (2005).

66. M. Stanke, O. Keller, I. Gunduz, A. Hayes, S. Waack, B. Morgenstern, AUGUSTUS: ab initio prediction of alternative transcripts. Nucleic Acids Res. 34, W435–9 (2006).

67. J. T. Lovell, A. Sreedasyam, M. E. Schranz, M. Wilson, J. W. Carlson, A. Harkess, D. Emms, D. M. Goodstein, J. Schmutz, GENESPACE tracks regions of interest and gene copy number variation across multiple genomes. Elife 11 (2022).

68. R Core Team, “R: A language and environment for statistical computing” (R Foundation for Statistical Computing, Vienna, Austria, 2024); https://www.R-project.org/.

69. D. M. Emms, S. Kelly, OrthoFinder: phylogenetic orthology inference for comparative genomics. Genome Biol. 20, 238 (2019).

70. Y. Wang, H. Tang, J. D. Debarry, X. Tan, J. Li, X. Wang, T.-H. Lee, H. Jin, B. Marler, H. Guo, J. C. Kissinger, A. H. Paterson, MCScanX: a toolkit for detection and evolutionary analysis of gene synteny and collinearity. Nucleic Acids Res. 40, e49 (2012).

71. H. Li, Minimap2: pairwise alignment for nucleotide sequences. Bioinformatics 34, 3094–3100 (2018).

72. D. J. Roiger, S. N. Jeffers, Evaluation of *Trichoderma spp.* for biological control of *Phytophthora* crown and root rot of apple seedlings. Phytopathology 81, 910–917 (1991).

73. B. Holmes, Evaluation of *Phytophthora parasitica* var. *nicotianae* for biocontrol of *Phytophthora parasitica* on *Catharanthus roseus*. Plant Dis 78, 193–199 (1994).

74. S. Jeffers, J. B. James, P. Sisco, E. Goheen, S. Frankel, Screening for resistance to *Phytophthora cinnamomi* in hybrid seedlings of American chestnut. 188–194 (2009).

75. J. Catchen, P. A. Hohenlohe, S. Bassham, A. Amores, W. A. Cresko, Stacks: an analysis tool set for population genomics. Mol. Ecol. 22, 3124–3140 (2013).

76. H. Li, R. Durbin, Fast and accurate short read alignment with Burrows-Wheeler transform. Bioinformatics 25, 1754–1760 (2009).

77. P. Danecek, J. K. Bonfield, J. Liddle, J. Marshall, V. Ohan, M. O. Pollard, A. Whitwham, T. Keane, S. A. McCarthy, R. M. Davies, H. Li, Twelve years of SAMtools and BCFtools. Gigascience 10 (2021).

78. G. A. Van der Auwera, B. D. O’Connor, Genomics in the Cloud: Using Docker, GATK, and WDL in Terra (“O’Reilly Media, Inc.,” 2020).

79. R. Poplin, V. Ruano-Rubio, M. A. DePristo, T. J. Fennell, M. O. Carneiro, G. A. Van der Auwera, D. E. Kling, L. D. Gauthier, A. Levy-Moonshine, D. Roazen, K. Shakir, J. Thibault, S. Chandran, C. Whelan, M. Lek, S. Gabriel, M. J. Daly, B. Neale, D. G. MacArthur, E. Banks, Scaling accurate genetic variant discovery to tens of thousands of samples, bioRxiv (2018)p. 201178.

80. B. L. Browning, X. Tian, Y. Zhou, S. R. Browning, Fast two-stage phasing of large-scale sequence data. Am. J. Hum. Genet. 108, 1880–1890 (2021).

81. B. L. Browning, Y. Zhou, S. R. Browning, A one-penny imputed genome from next-generation reference panels. Am. J. Hum. Genet. 103, 338–348 (2018).

82. R. Corbett-Detig, R. Nielsen, A Hidden Markov Model Approach for Simultaneously Estimating Local Ancestry and Admixture Time Using Next Generation Sequence Data in Samples of Arbitrary Ploidy. PLoS Genet. 13, e1006529 (2017).

83. A. M. Sandercock, J. W. Westbrook, Q. Zhang, H. A. Johnson, T. M. Saielli, J. A. Scrivani, S. F. Fitzsimmons, K. Collins, M. T. Perkins, J. H. Craddock, J. Schmutz, J. Grimwood, J. A. Holliday, Frozen in time: Rangewide genomic diversity, structure, and demographic history of relict American chestnut populations. Mol. Ecol. 31, 4640–4655 (2022).

84. J. Goudet, hierfstat, a package for r to compute and test hierarchical *F*-statistics. Mol. Ecol. Notes 5, 184–186 (2005).

85. K. W. Broman, H. Wu, S. Sen, G. A. Churchill, R/qtl: QTL mapping in experimental crosses. Bioinformatics 19, 889–890 (2003).

86. D. Grattapaglia, R. Sederoff, Genetic linkage maps of Eucalyptus grandis and Eucalyptus urophylla using a pseudo-testcross: mapping strategy and RAPD markers. Genetics 137, 1121–1137 (1994).

87. D. H. Alexander, K. Lange, Enhancements to the ADMIXTURE algorithm for individual ancestry estimation. BMC Bioinformatics 12, 246 (2011).

88. T. J. Ellis, D. L. Field, N. H. Barton, Efficient inference of paternity and sibship inference given known maternity via hierarchical clustering. Mol. Ecol. Resour., doi: 10.1111/1755-0998.12782 (2018).

89. D. G. Butler, B. R. Cullis, A. R. Gilmour, B. J. Gogel, R. Thompson, ASReml-R reference manual version 4. VSN International Ltd, Hemel Hempstead, HP1 1ES, UK (2017).

90. P. H. Noah, N. L. Cagle, J. W. Westbrook, S. F. Fitzsimmons, Identifying resilient restoration targets: Mapping and forecasting habitat suitability for *Castanea dentata* in Eastern USA under different climate-change scenarios. Climate Change Ecology 2, 100037 (2021).

91. R. J. Hijmans, J. Van Etten, M. Mattiuzzi, M. Sumner, J. A. Greenberg, O. P. Lamigueiro, A. Bevan, E. B. Racine, A. Shortridge, Raster package in R. Version. https://Mirro Rs. Sjtug. Sjtu. Edu. Cn/Cran/Web/Packa Ges/Rast er/Raster. Pdf (2013).

92. L. H. Moro Rosso, A. F. de Borja Reis, A. A. Correndo, I. A. Ciampitti, XPolaris: an R-package to retrieve United States soil data at 30-meter resolution. BMC Res. Notes 14, 327 (2021).

93. O. F. Christensen, M. S. Lund, Genomic prediction when some animals are not genotyped. Genet. Sel. Evol. 42, 2 (2010).

94. I. Aguilar, I. Misztal, D. L. Johnson, A. Legarra, S. Tsuruta, T. J. Lawlor, Hot topic: a unified approach to utilize phenotypic, full pedigree, and genomic information for genetic evaluation of Holstein final score. J. Dairy Sci. 93, 743–752 (2010).

95. A. Legarra, O. F. Christensen, I. Aguilar, I. Misztal, Single Step, a general approach for genomic selection. Livest. Sci. 166, 54–65 (2014).

96. O. F. Christensen, P. Madsen, B. Nielsen, T. Ostersen, G. Su, Single-step methods for genomic evaluation in pigs. Animal 6, 1565–1571 (2012).

97. J. Ødegård, T. H. E. Meuwissen, Estimation of heritability from limited family data using genome-wide identity-by-descent sharing. Genet. Sel. Evol. 44, 16 (2012).

98. S. Gezan, D. Murray, A. A. de Oliveira, G. Galli, VSN International, ASRgenomics: Complementary Genomic Functions. [Preprint] (2024). https://CRAN.R-project.org/package=ASRgenomics.

99. P. M. VanRaden, Efficient methods to compute genomic predictions. J. Dairy Sci. 91, 4414–4423 (2008).

100. P. C. Austin, J. Merlo, Intermediate and advanced topics in multilevel logistic regression analysis. Stat. Med. 36, 3257–3277 (2017).

101. A. Dahl, V. Iotchkova, A. Baud, Å. Johansson, U. Gyllensten, N. Soranzo, R. Mott, A. Kranis, J. Marchini, A multiple-phenotype imputation method for genetic studies. Nat. Genet. 48, 466–472 (2016).

102. H. D. Daetwyler, M. P. L. Calus, R. Pong-Wong, G. de Los Campos, J. M. Hickey, Genomic prediction in animals and plants: simulation of data, validation, reporting, and benchmarking. Genetics 193, 347–365 (2013).

103. J. W. R. Martini, M. F. Schrauf, C. A. Garcia-Baccino, E. C. G. Pimentel, S. Munilla, A. Rogberg-Muñoz, R. J. C. Cantet, C. Reimer, N. Gao, V. Wimmer, H. Simianer, The effect of the H-1 scaling factors τ and ω on the structure of H in the single-step procedure. Genet. Sel. Evol. 50, 16 (2018).

104. M. L. Cipollini, J. P. Moss, W. Walker, N. Bailey, C. Foster, H. Reece, C. Jennings, Evaluation of an Alternative Small Stem Assay for Blight Resistance in American, Chinese, and Hybrid Chestnuts (*Castanea* spp.). Plant Dis. 105, 576–584 (2021).

105. C. E. Conn, N. Howie, M. Lynch, S. Lee, E. Young, J. Westbrook, J. Holliday, Q. Zhang, M. L. Cipollini, Validation of an alternative small stem assay for blight resistance in chestnut seedlings and recommendations for broader use. Plant Dis. 107, 1576–1583 (2023).

106. M. R. Robinson, A. W. Santure, I. Decauwer, B. C. Sheldon, J. Slate, Partitioning of genetic variation across the genome using multimarker methods in a wild bird population. Mol. Ecol. 22, 3963–3980 (2013).

107. P. Medina, B. Thornlow, R. Nielsen, R. Corbett-Detig, Estimating the Timing of Multiple Admixture Pulses During Local Ancestry Inference. Genetics 210, 1089–1107 (2018).

108. J. T. Lovell, qtlTools: Data Processing and Plotting in Association with R/qtl. R package version.

109. K. W. Broman, D. M. Gatti, P. Simecek, N. A. Furlotte, P. Prins, Ś. Sen, B. S. Yandell, G. A. Churchill, R/qtl2: Software for mapping quantitative trait loci with high-dimensional data and multiparent populations. Genetics 211, 495–502 (2019).

110. J. Yang, N. A. Zaitlen, M. E. Goddard, P. M. Visscher, A. L. Price, Advantages and pitfalls in the application of mixed-model association methods. Nat. Genet. 46, 100–106 (2014).

111. Broman, K. W., & Sen, Ś., A Guide to QTL Mapping with R/qtl (2009). https://rqtl.org/book/rqtlbook_ch04.pdf.

112. R. C. Gaynor, G. Gorjanc, J. M. Hickey, AlphaSimR: an R package for breeding program simulations. G3 11 (2021).

113. S. Chang, J. Puryear, J. Cairney, A simple and efficient method for isolating RNA from pine trees. Plant Mol. Biol. Rep. 11, 113–116 (1993).

114. A. Dobin, C. A. Davis, F. Schlesinger, J. Drenkow, C. Zaleski, S. Jha, P. Batut, M. Chaisson, T. R. Gingeras, STAR: ultrafast universal RNA-seq aligner. Bioinformatics 29, 15–21 (2013).

115. S. Chen, Y. Zhou, Y. Chen, J. Gu, fastp: an ultra-fast all-in-one FASTQ preprocessor. Bioinformatics 34, i884–i890 (2018).

116. Y. Liao, G. K. Smyth, W. Shi, The Subread aligner: fast, accurate and scalable read mapping by seed-and-vote. Nucleic Acids Res. 41, e108 (2013).

117. M. Love, S. Anders, W. Huber, Differential analysis of count data – the DESeq2 package. Genome Biol. (2013).

118. M. Dowle, A. Srinivasan, data.table: Extension of ‘data.framè. (2023). https://CRAN.R-project.org/package=data.table.

119. H. Pagès’, P. Aboyoun, R. Gentleman, S. DebRoy, Biostrings: Efficient Manipulation of Biological Strings (2024; https://bioconductor.org/packages/Biostrings).

120. P. Li, X. Quan, G. Jia, J. Xiao, S. Cloutier, F. M. You, RGAugury: a pipeline for genome-wide prediction of resistance gene analogs (RGAs) in plants. BMC Genomics 17, 852 (2016).

121. J. Sperschneider, P. N. Dodds, EffectorP 3.0: Prediction of apoplastic and cytoplasmic effectors in fungi and oomycetes. Mol. Plant. Microbe. Interact. 35, 146–156 (2022).

122. J. A. Crouch, A. Dawe, A. Aerts, K. Barry, A. C. L. Churchill, J. Grimwood, B. I. Hillman, M. G. Milgroom, J. Pangilinan, M. Smith, A. Salamov, J. Schmutz, J. S. Yadav, I. V. Grigoriev, D. L. Nuss, Genome sequence of the chestnut blight fungus *Cryphonectria parasitica* EP155: A fundamental resource for an archetypical invasive plant pathogen. Phytopathology 110, 1180–1188 (2020).

123. F. Teufel, J. J. Almagro Armenteros, A. R. Johansen, M. H. Gíslason, S. I. Pihl, K. D. Tsirigos, O. Winther, S. Brunak, G. von Heijne, H. Nielsen, SignalP 6.0 predicts all five types of signal peptides using protein language models. Nat. Biotechnol. 40, 1023–1025 (2022).

124. D. E. Beck, Department of Agriculture, Forest Service, Southeastern Forest Experiment Station (Asheville, NC: U.S, 1962)vol. 2.

